# Feedback-related EEG dynamics separately reflect decision parameters, biases, and future choices

**DOI:** 10.1101/2021.05.10.443374

**Authors:** Hans Kirschner, Adrian G. Fischer, Markus Ullsperger

## Abstract

Optimal decision making in complex environments requires dynamic learning from unexpected events. To speed up learning, we should heavily weight information that indicates state-action-outcome contingency changes and ignore uninformative fluctuations in the environment. Often, however, unrelated information is hard to ignore and can potentially bias our learning. Here we used computational modelling and EEG to investigate learning behaviour in a modified probabilistic choice task that introduced two types of unexpected events that were irrelevant for optimal task performance, but nevertheless could potentially bias learning: pay-out magnitudes were varied randomly and, occasionally, feedback presentation was enhanced by visual surprise. We found that participants’ overall good learning performance was biased by distinct effects of these non-normative factors. On the neural level, these parameters are represented in a dynamic and spatiotemporally dissociable sequence of EEG activity. Later in feedback processing the different streams converged on a central to centroparietal positivity reflecting a final pathway of adaptation that governs future behaviour.

## Introduction

Learning to accurately predict desirable or undesirable outcomes based on beliefs about states of the world is crucial for adaptive behaviour. A critical challenge during learning is that most state-action-outcome contingencies fluctuate from one moment to the next and sometimes even reverse. Consider for example the food in your favourite restaurant. Even though the food quality might vary to some degree (e.g., mood of the chef, availability of fresh ingredients), you have learnt to revisit this place for a tasty meal. However, after the retirement of the head chef, you might disagree with the taste of the new chef and must search for a new reliable source for a delicious meal.

A wealth of studies report, that people can adaptively integrate new information into their beliefs about the world by considering the environments stochasticity and volatility (Behrens et al., 2007; d’Acremont & Bossaerts, 2016; Diederen et al., 2016; Nassar et al., 2019; Summerfield & Tsetsos, 2015). Specifically, learning in dynamic environments is driven by prediction errors (i.e., a simple form of surprise: is the outcome better or worse than expected and by how much?) and an individual learning-rate that scales the impact of prediction errors on value updates (Sutton & Barto, 2018). Recent evidence suggests that human participants are capable to adaptively calibrate the learning rate according to the statistical context of the environment, whereby prediction errors caused by uninformative outliers (oddballs) are down-weighted and prediction errors in volatile environments up-weighted (d’Acremont & Bossaerts, 2016; McGuire et al., 2014; Nassar et al., 2019). The P3b, a stimulus-locked centroparietal positivity in the EEG that has previously been associated with adjustments in learning behaviour (Jepma et al., 2018; Jepma et al., 2016; Polich, 2007), has been shown to be modulated by the current learning rate, both after factual and counterfactual outcome information (Fischer & Ullsperger, 2013). Indeed, a recent study explicitly showed, that the P3b reflects neuronal processes that play an important role in adaptively calibrating learning, depending on the statistical context of the prediction error (Nassar et al., 2019). In this study, the amplitude of the P3b positively predicted learning in the context of stimulus-outcome contingencies reversals, but negatively predicted learning in the presence of (uninformative) oddballs.

Importantly, in complex environments additional factors can influence learning, some of which even may be uninformative for upcoming decisions. Considering the restaurant example alluded to previously, one’s expectation about having a tasty meal at a particular restaurant could also be influenced by factors not directly related to the food quality. For example, being soaked by a sudden rainstorm on the way to the restaurant can spoil the whole evening and perhaps also worsen the memory of food quality despite an excellent meal. Indeed, there is evidence, that non-normative factors such as randomly varying outcome magnitudes are tied to learning and thus biasing future outcome expectations (Daw et al., 2011; Fischer et al., 2017). In a previous study, where participants had to catch bags dropped from a helicopter hidden in the clouds, they learned quickly to infer the helicopter’s location based on where the bags fell down (McGuire et al., 2014). Interestingly, participants updated their belief about the helicopter location more on rare trials, when the bags contained gold rather than stones. This bias in belief update could have been driven (a) by the higher obtained reward and/or (b) by the unexpectedness, i.e., salience of the gold bags. A major open question in learning and decision-making field is how the human brain integrates normative and biasing influences to guide future behaviour.

To address this question, we combined computational modelling, electroencephalography (EEG), and decoding to study distinct effects of normative and non-normative factors on learning. Specifically, we measured learning behaviour in a modified probabilistic reversal-learning task that introduced two types of non-normative factors that were irrelevant for optimal task performance, but nevertheless could potentially bias learning: pay-out magnitudes were varied randomly and, occasionally, feedback presentation was enhanced by visual surprise. We hypothesized that participants are unable to ignore these non-normative task factors despite explicit knowledge of their irrelevance for task performance. This should be reflected in biased learning, choice behaviour and independent feedback-induced EEG dynamics. In line with our expectations, we demonstrate that participants’ overall good learning performance was indeed biased by random pay-out magnitudes and sensory surprise. On the behavioural level, these biases manifest themselves by slowing on consecutive trials and in future decisions on the same stimuli. Modelling results show that participants’ learning was influenced by two biases: (1) they reweighted the outcomes depending on the (irrelevant) magnitude and (2) were distracted by visual surprise to an individually variable degree. On the neural level, these biases were accompanied by distinct feedback-locked EEG correlates that predict behavioural adaptations.

## Results

We measured surprise and learning signals in the EEG of twenty-eight human participants while they performed a reversal learning variant of an established probabilistic learning task (Fischer & Ullsperger, 2013) that was specifically tailored to our research question. On each trial of the task, participants had to choose or avoid gambling on a centrally presented stimulus (see Figure 1Ai and Methods for additional information on the task). Gambling on a stimulus resulted in a monetary gain or loss depending on the reward probability (20%, 50% or 80%) of the respective stimuli. If subjects decided not to gamble, they avoided any financial consequences, but were still able to observe what would have happened if they had gambled. Hence, two types of learning could occur: learning from real rewarding or punishing outcomes, or from purely informative, counterfactual outcomes without direct reward or punishment. During the task, reward probabilities of the stimuli could change unexpectedly, such that a good stimulus, that would usually lead to gain of money, could change to be either neutral (random pay-outs), or bad and vice versa. Moreover, we introduced two types of unexpected events in the task: a) simple visual surprise, where on some trials, the colour of the feedback background changed to match the feedback colour (see Figure 1Aii), and b) pay-out magnitudes randomly varying between either 10 or 80 points (see Figure 1Aiii). Importantly, we explicitly informed the participants, that pay-out magnitudes varied randomly and therefore should be ignored. We did not brief participants about the occasionally changing feedback background.

In general, participants’ choices followed the reward probabilities of the stimuli (see Figure 1Bii and Supplementary Figure 4), suggesting that the participants learned the task well. In addition, we observed no difference in the absolute number of correct decisions following good as compared to bad stimuli (*t*(23) = 1.31, *p* = 0.20). This suggests that participants learnt equally well from factual and counterfactual outcomes.

### Visual surprise and pay-out magnitudes modulate participants’ behaviour

To investigate the factors influencing or biasing decisions, we performed regression analyses, predicting either participants’ decisions to choose or avoid gambling, or their reaction times (RT) on a given trial (see Method section and Supplementary Results for a detailed description of the models and results). To study immediate trial effects we submitted the original trial order, where the three stimuli were intermixed. Neither preceding visual surprise (bGLM1; previous flash: *t*(23) = −.52, *p*(corrected) = 1) nor higher previous outcome magnitudes (bGLM1; previous magnitude: *t*(23) = −1.21, *p*(corrected) = 1) affected participants’ choices (see Figure 2Ai). Interestingly, we found that both types of unexpected events had a slowing effect on RTs on subsequent trials (see Figure 2Aii; bGLM2; previous flash: ΔRT = ∼23ms; *t*(23) = 4.23, *p*(corrected) = .002; previous magnitude: ΔRT = ∼12ms; *t*(23) = 3.32, *p*(corrected) = .029).

Learning was investigated in a separate model (see methods and Figure2B). We demonstrate, that choosing to gamble on a stimulus became more likely after previous wins (bGLM3; previous outcome: *t*(23) = 16.95, *p*(corrected) < .001) and when the stimulus was chosen before (bGLM3; previous choice: *t*(23) = 13.91, *p*(corrected) < .001). Importantly, these effects were modulated by the previous reward magnitude (bGLM3; previous magnitude x previous outcome: *t*(23) = 5.61, *p*(corrected) < .001; previous magnitude x previous outcome x previous choice: *t*(23) = −2.83, *p*(corrected) = .06). Follow-up analyses revealed that participants were more likely to gamble on a stimulus after wins with high reward magnitudes (*t*(23) = 5.71, *p* < .001, d = 0.49) and less likely to choose a stimulus after high losses (*t*(23) = 2.82, *p* = 0.01, d = 0.21). Moreover, subjects were more likely to gamble on a stimulus after having received a high counterfactual reward (*t*(23) = 6.42, p < .001, d = 0.76). On the contrary, participants were less likely to choose a stimulus after high factual losses (*t*(23) = 2.62, *p* = 0.02, d = 0.42). These results may reflect an enhancement of simple win-stay/lose-shift heuristics (Nowak & Sigmund, 1993) during decision making by high reward magnitudes. Receiving visually surprising feedback during a previous stimulus encounter did not influence participants’ choices or switch probability (Supplementary Figure 5B and 6). In addition, there was no systematic modulation of participants’ RTs through previous choice, outcome, magnitude, or feedback background colour of the same stimulus beyond the effect of immediately preceding outcomes (bGLM4, Figure 2).

**Figure 1.**
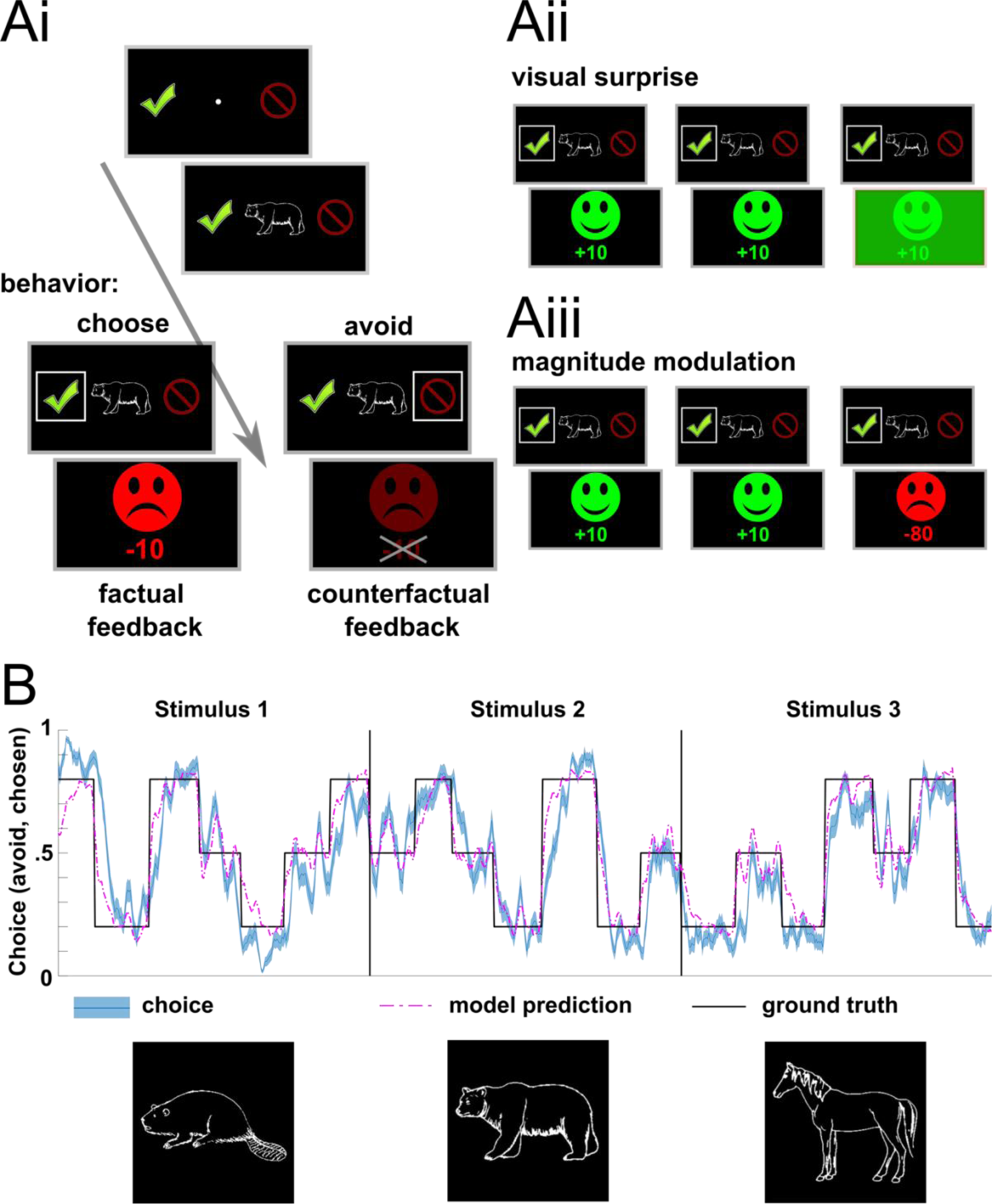
Experimental design, observed and modelled behaviour. **(Ai)** Time course of the task. On each trial, a fixation dot and choice options were presented for 300 – 700 ms. Next, the stimulus was presented for up to 1700 ms. During this time participants had to decide, if they want to gamble on the stimulus or not. After subject’s decision, their choice was highlighted for 350 ms. Finally, depending on participants choice, either factual or counterfactual feedback was presented for 750 ms. **(Aii)** On 20% of the trials, the colour of the feedback background changed from black to a feedback-matching colour (i.e. red or green), introducing visual surprise to the task. **(Aiii)** A second manipulation of surprise in the task focused on the reward magnitudes. Here, magnitudes randomly varied between 10 and 80 points. **(B)** Modelled and choice behaviour of the participants in the task, stretched out for all stimuli. Ground truth represents the current reward probability (20%, 50% or 80%) of the respective stimulus.

**Figure 2.**
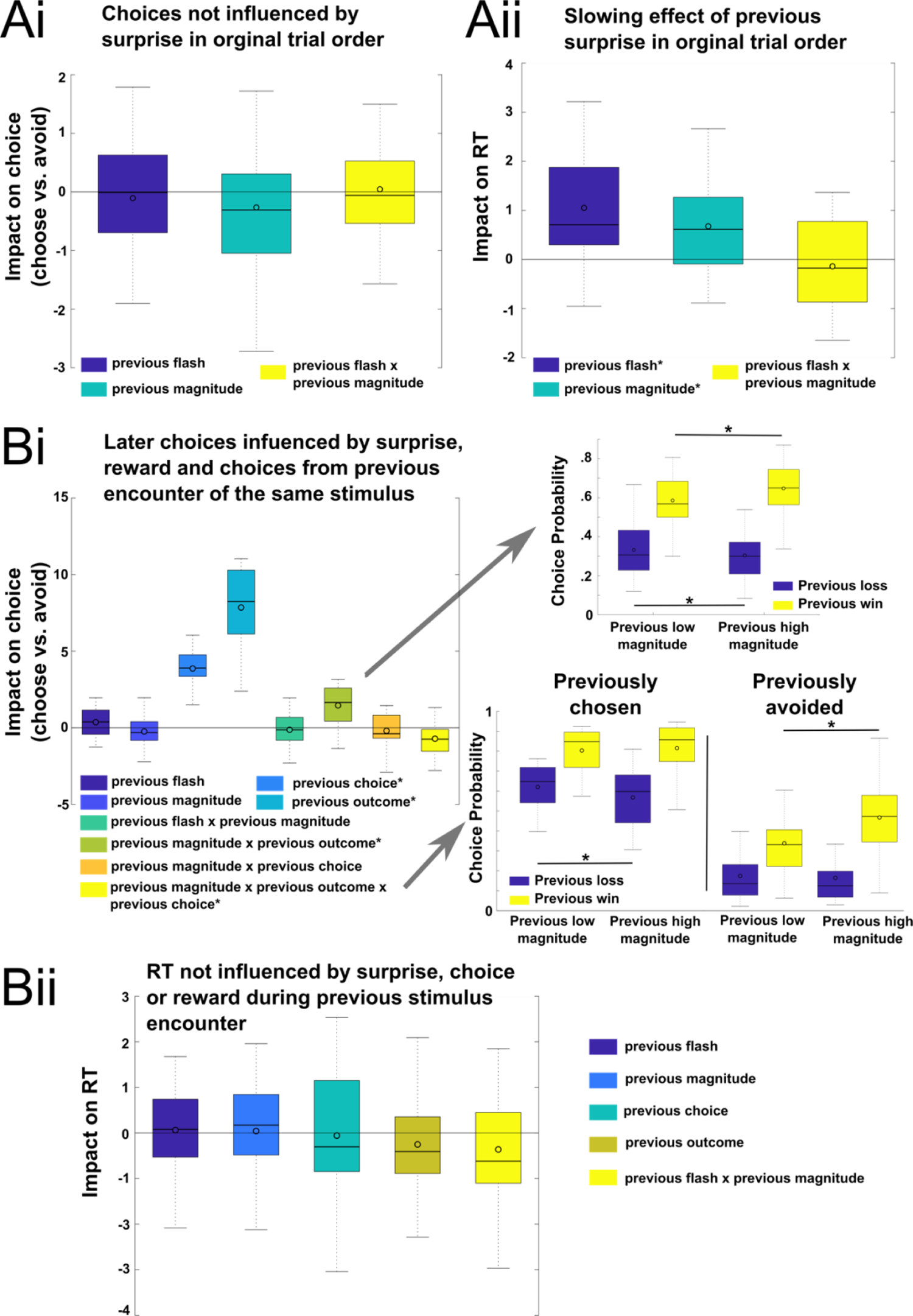
Behavioural Results. **(A)** Main results for regressions on original trial order, investigating the impact of visual surprise and outcome magnitude on choices **(Ai)** and RT **(Aii)** on the next trial while ignoring stimulus identity. We found that visual surprise and high outcome magnitude slowed down subsequent responses but did not affect participants’ choices. **(B)** Regression analyses on a resorted trial order according to appearance of the same stimulus identity show that reward magnitudes influenced participants decisions **(Bi)** but not their RTs **(Bii)**. *Note:* Impact is quantified and visualized as averaged within-subject *t*-values. *Significant regressor (derived from t-tests of the individual regression t-values against zero). Boxes = interquartile range (IQR), − = median, o = mean, whiskers =1.5 × IQR.

Taken together, both, random pay-out magnitudes and sensory surprise have an impact on behaviour in the task. Specifically, we show that visual surprise and random reward magnitudes are similarly processed, in that they both increase RT on consecutive trials. This potentially demonstrates a comparable and non-specific surprise evaluation. Moreover, we demonstrate that non-normative task factors bias learning, despite explicit knowledge of its irrelevance. Critically, it is not possible to disambiguate which processes drive this learning bias within these analyses. We therefore utilized computational modelling to quantify potential variables involved in this learning bias.

### Unexpected, non-normative factors in the task bias learning by scaling the size and impact of RPE

We fitted four reinforcement learning models (see methods for a detailed description of computational modelling procedure) to participants’ behaviour. The optimal way to maximize outcome in this task, is to ignore random pay-out magnitudes and visual surprise during decision making. However, we hypothesized that participants are unable to ignore non-normative factors in the task. Indeed, modelling results suggest that participants’ learning was influenced by two biases: (1) they reweighted the outcomes depending on the (actually irrelevant) magnitude and (2) were distracted by the visual surprise to an individually variable degree. In the best fitting model, this was operationalized by a reward-scaling parameter gamma (γ< .5 indicated up-weighting of low outcome magnitudes; γ> .5 indicated down-weighting of high outcome magnitudes; Methods) and an attention parameter lambda which scaled the individual learning rate on trials with visual surprise (λ < 0 suggests down-weighting and λ > 0 up-weighting of the individual learning rate).

In our sample, the average reward-scaling parameter gamma was smaller than the indifference value (γ = .5, where high and low magnitudes outcomes have identical reward magnitudes; *t*(23) = −10.77, *p* < .001, d = −2.25, CI [0.14, 0.26]), suggesting that participants down-weighted low magnitude outcome leading to decreased value updating (Figure 3). Interestingly, we found that the less a participant’s choices were biased by magnitudes (the larger, i.e., the closer gamma was to .5), the better was the participant’s performance in the task (Figure 3C). The lambda parameter was significantly smaller than 0 (*t*(23) = −2.54, *p* = 0.02, d = −0.53, CI [−0.28, −0.03], see Figure 3B), suggesting that sensory surprise did not get ignored, but had a rather distracting effect by reducing the learning rate (and thereby down-weighting the effect of the reward prediction error on value update) on respective trials. The learning bias induced by visual surprise reflected in lambda did not correlate with overall task performance (Figure 3C). Interestingly, when examining the distribution of the lambda parameter, participants appear to cluster into two groups: Individuals distracted by visual surprise (i.e., they update their values less on trials with visual surprise; bright dots in Figure 3C), and individuals with small positive lambda scores, suggesting slightly enhanced learning through increased feedback salience by visual surprise (dark dots in Figure 3C). In addition, there was a trend for a positive correlation between the absolute difference to the indifference value between gamma and lambda scores (*r* = .20, *p* = .08), suggesting a general susceptibility to learning biases that would be interesting to follow up in larger samples.

**Figure 3.**
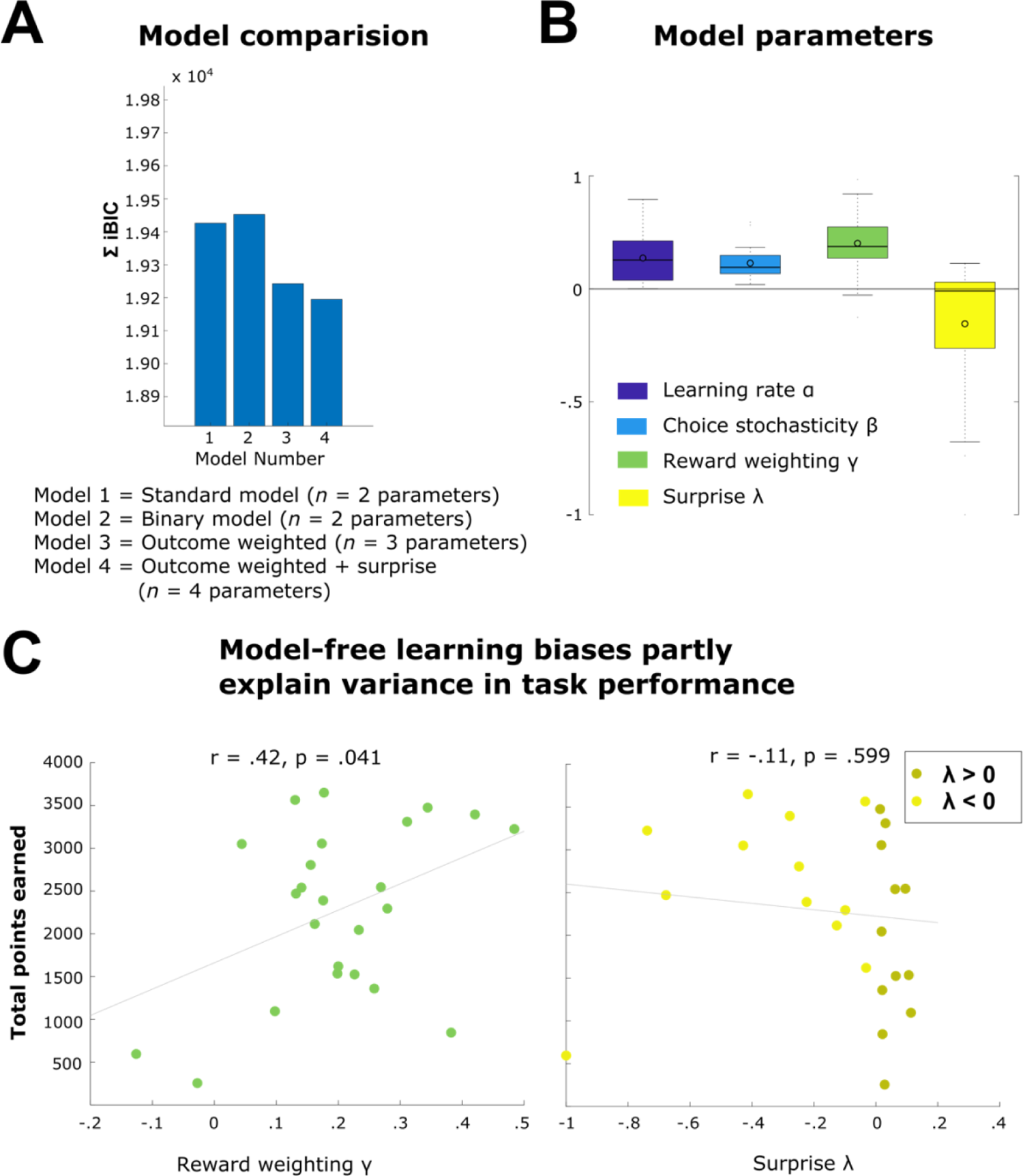
Computational modelling results. **(A)** Comparison of the candidate models by iBIC indicates, that model 4 that allows for free reward scaling and the learning rate to be changed by surprise fit the data best. **(B)** Fitted parameter distribution of model 4. Boxes = interquartile range (IQR), − = median, o = mean, whiskers =1.5 × IQR. **(C)** depicts the correlation between model parameters reflecting learning biases and task performance.

Model comparison revealed, that neither the alternative model that weighted high outcomes eight times as important as low outcomes (standard RL model), nor the model that weighted outcomes equally (binary model, the optimal way to learn in the task), fit participants behaviour better than the model accounting for outcome reweighting and visual surprise provided the best fit to the data (Figures 3, 1Bii). This was supported by model and parameter recovery analyses (Supplementary Figure 1).

Taken together, the behavioural results suggest that the non-normative factors in the task affected participants’ behaviour. Both random pay-out magnitudes and sensory surprise induce learning biases. These learning biases in part explain the variance in task performance between participants.

### Neural correlates

We submitted feedback-locked EEG epochs to mass-univariate multiple robust regression analysis across single trials to reveal representations of the variables driving or biasing value update and future choice behaviour, namely reward prediction errors (RPE), outcome magnitudes, and visual surprise.

### EEG correlates of reward prediction errors are dynamic and differ between real and fictive outcomes

Replicating previous findings (Fischer & Ullsperger, 2013), we found an early dissociation of feedback processing between real and fictive outcomes. First, an occipital negative early reward prediction error (RPE) effect that occurred 180-220 ms after feedback onset was exclusively significant for fictive feedback (peak at Oz 210 ms, *t*(23) = −4.99, *p* < .001, d = −1.04, CI [−2.42, −1.00]). Second, only real outcomes were associated with an early frontal positive prediction error effect spanning from 200-290 ms (peak at FCz 260 ms, *t*(23) = 5.07, *p* < .001, d = 1.06, CI [1.21, 2.88]) and a subsequent negative frontal prediction error covariation in the time range of 330-400 ms with a peak at electrode FCz at 360 ms, *t*(23) = − 4.47, *p* < .001, d = −0.93, CI [−3.46, −1.27], see Figure 4A). As depicted in Figure 5A, in the averaged event-related potentials these covariations reflect the feedback-related negativity (FRN, Walsh & Anderson, 2012) and P3a components (see Figure5A). Formal tests of the exclusiveness of these effects to real and fictive feedback processing are reported in Supplementary Figure 7. Similar to Fischer & Ullsperger (2013), we find that, as feedback processing continues, real and fictive outcomes converge on a similar parietal RPE correlate that matches the P3b ERP component (real, peak at Pz 560 ms, *t*(23) = −5.09, *p* < .001, d = − 1.06, CI [−2.50, −1.06]; fictive, peak at Pz 430 ms, *t*(23) = −4.99, *p* < .001, d = −1.04, CI [−2.42, −1.00]; see Figure 4A and Figure 5A). The directions of the parietal RPE effects are such that for both, real and fictive outcomes, the ERP is more positive-going for more unfavourable outcomes (i.e. more negative RPEs when the gamble was chosen and more positive RPEs when the gamble was avoided).

**Figure 4.**
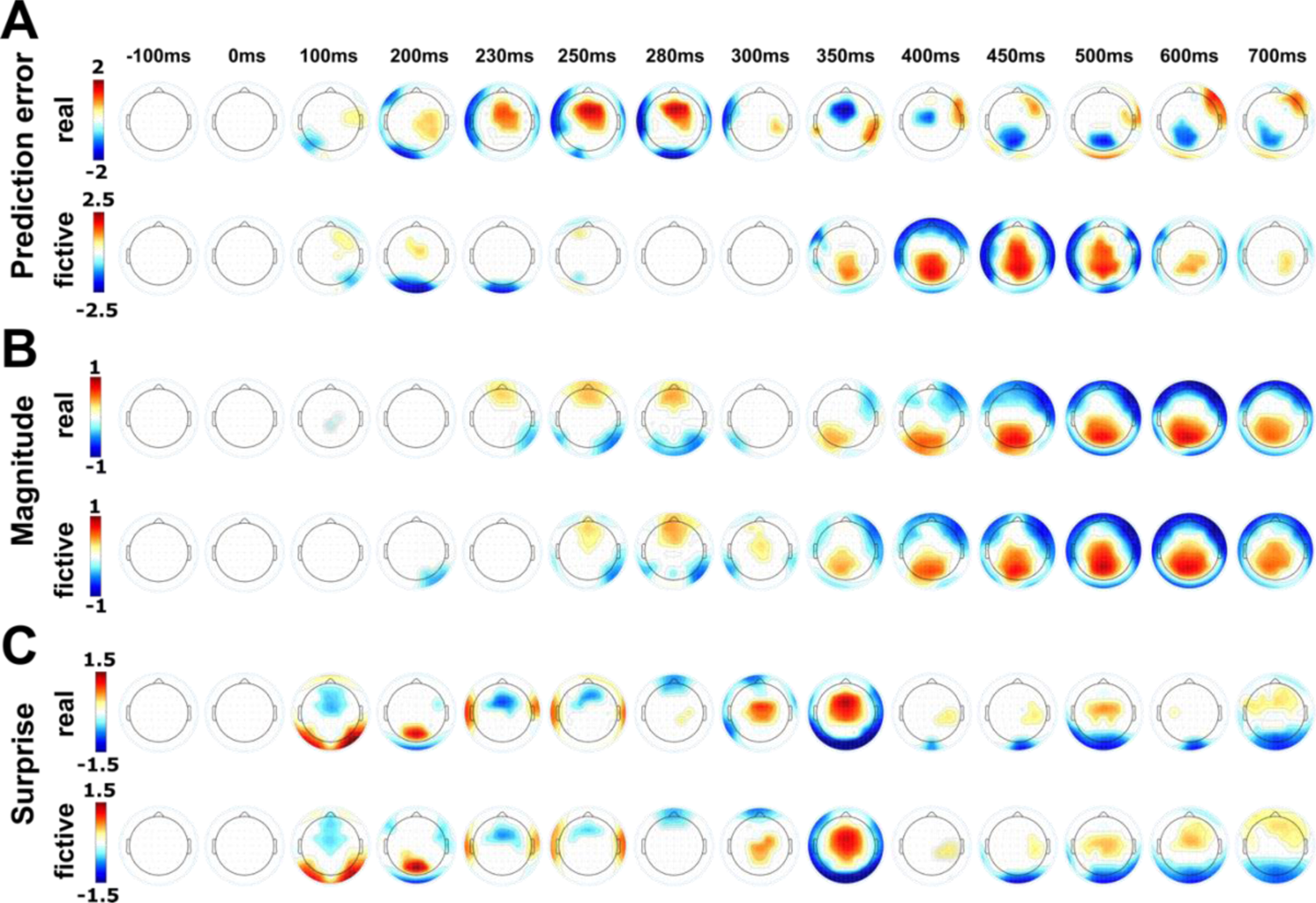
Single-trial regression analysis of feedback-locked EEG epochs. Time course of regression weight topographies for the feedback-locked reward prediction error **(A)**, magnitude **(B),** and surprise **(C)** regressor split up into real and fictive outcomes. Topographies show beta coefficients thresholded at critical p-value from FDR correction.

**Figure 5.**
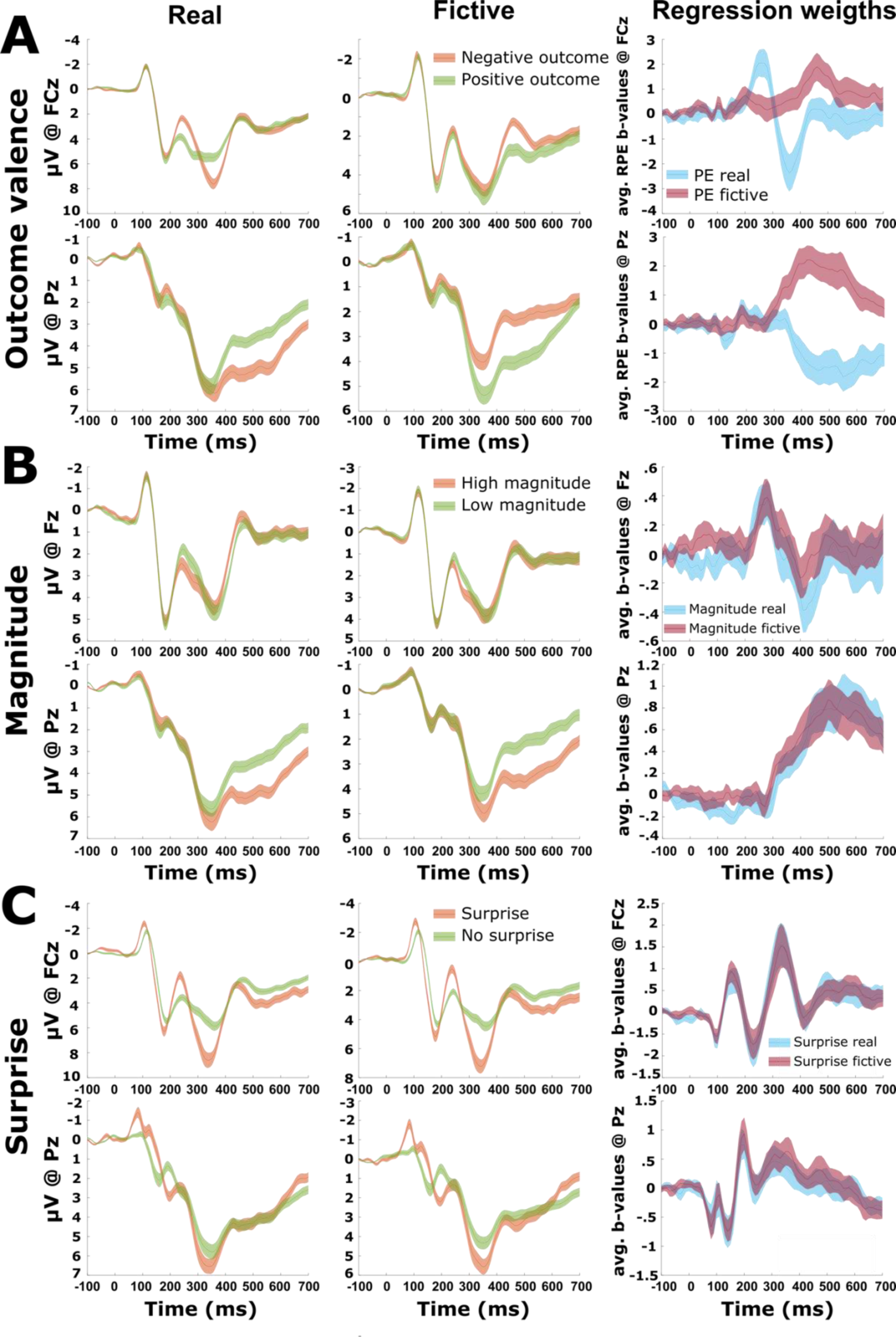
EEG representations of outcome parameters reward prediction error, magnitude, and surprise contribute differentially to feedback-locked event-related EEG dynamics and ERPs. The first two columns show feedback-locked event-related potential (ERP) waveforms separately for real and fictive feedback split up for the main task manipulations outcome (A), magnitude (B) and visual surprise (C). The column on the right depicts the regression time courses for the according main effects in the models eegGLM 1 & 2, comparing processing of real (blue) and fictive (purple) feedback. Shown are electrodes of maximal effects of early (FRN, P3a at FCz) and late (P3b at Pz) ERP correlates. Shadings indicate 99% confidence intervals.

### Decision biases magnitude and surprise differentially modulate feedback-related ERPs

As can be seen in Figure 4B and 5B, reward magnitude is coded in a fairly similar fashion in real and fictive outcomes. Specifically, we found a positive frontal-to-frontocentral covariation in the time range of the FRN for both real (230 – 290 ms, peak at Fz 280 ms, *t*(23) = 3.88, *p* < .001, d = 0.81, CI [0.17, 0.57]) and fictive (260 – 290 ms, peak at Fz 280 ms, *t*(23) = 4.35, *p* < .001, d = 0.91, CI [0.20, 0.57]) feedback. In other words, larger outcome magnitudes are associated with more positive FRN amplitudes. Later in feedback processing we find a positive parietal covariation between the magnitude regressor and neural activity in the P3b range (real feedback: 320 – 790 ms, peak at Pz 550 ms, *t*(23) = 7.00, *p* < .001, d = 1.46, CI [0.65, 1.19]; fictive feedback: 310 – 750 ms, peat at Pz 500 ms, *t*(23) = 6.64, *p* < .001, d = 1.38, CI [0.60, 1.15]).

Neural correlates of visual surprise are very dynamic, yet similar in real and fictive feedback (see Figure 4C and 5C). First, there is an early frontocentral negative covariation combined with a positive bilateral occipital covariation around 100ms (real feedback: 70 – 110 ms, peak at FCz 90 ms, *t*(23) = −5.56, *p* < .001, d = −1.16, CI [−0.89, −0.41]; fictive feedback: *t*(23) = −4.67, *p* < .001, d = −0.97, CI [−0.77, −0.30]) followed by a positive covariation around 160 ms (real feedback: 130 −170, – 100 ms, peak at FCz 160 ms, *t*(23) = 3.48, p = 0.002, d = 0.72, CI [0.29, 1.16]; fictive feedback: *t*(23) = 5.27, *p* < .001, d = 1.10, CI [0.57, 1.31]). Next, we found a positive parieto-occipital covariation with polarity reversal at Oz around 200 ms (real feedback: 190 – 210, peak at Pz 200ms, *t*(23) = 4.48, *p* < .001, d = 0.95, CI [0.40, 1.09]; fictive feedback: *t*(23) = 4.74, *p* < .001, d = 0.98, CI [0.53, 1.35]). This was followed by a frontocentral negative (real feedback: 220 – 250 ms, peak at FCz 230 ms, *t*(23) = −4.39, *p* < .001, d = −0.92, CI [−1.40, −0.51]; fictive feedback: *t*(23) = −3.01, *p* = 0.006, d = −0.63, CI [−1.25, −0.23]) and a subsequent mid-latency strong positive covariation (real feedback: 290 – 390 ms, peak at FCz 330ms, *t*(23) = 4.46, *p* < .001, d = 0.93, CI [0.84, 2.28]; fictive feedback: *t*(23) = 4.92, *p* < .001, d = 1.03, CI [0.79, 1.94]), which contribute to the FRN and P3a, respectively (Figure 5C). Finally, there was a small central positive covariation contributing to the P3b with a shorter time window in the real (490 – 510 ms, peak at Cz 500ms, *t*(23) = 3.34, *p* = 0.003, d = 0.70, CI [0.25, 1.05]) as compared to the fictive feedback condition (470 – 610 ms, peak at Pz 590 ms, *t*(23) = 2.87, *p* = 0.009, d = 0.60, CI [0.15, 0.93]).

Taken together, the results from eegGLM1 and eegGLM2 revealed that reward prediction errors are initially processed differently in real and fictive outcomes but converge on final neural pathway reflected in similar EEG dynamics for both conditions. This suggests that learning from fictive feedback engages a specific neural mechanism distinct from learning from real outcomes. In contrast, neural correlates of magnitude and visual surprise effects are similar for real and fictive outcomes and broadly overlap with RPE processing in real outcomes temporally and spatially. Interestingly, these results suggest, that both magnitude and surprise effects contribute to the FRN/P3 ERP complex over and above the effects of reward prediction errors.

### Feedback-related EEG dynamics predict future behaviour

Next, we investigated the relationship between EEG activity during feedback processing and behavioural switches on the next encounter of stimuli with the same identity. We used multivariate pattern analysis on ERP activity of the whole scalp and trained a support vector machine to predict behavioural switches. As can be seen in Figure 6A, cluster-based permutation analyses revealed a large cluster of time points in which the decoding of future switches was significantly greater than chance level. Interestingly, this time window is spanning over all the significant neural correlates of the regressors of interest in eegGLM 1&2, ramping up from 110 ms across the FRN/P3a latency range, plateauing at its maximum in the P3b time window, and tapering off until 790 ms after feedback presentation. A search light analysis at the maximal individual decoding accuracy revealed a topographic match with the P3b which occurs in this time range (see Figure 6 Bi). To test whether the outcome parameters represented in the feedback-locked EEG dynamics contribute to the EEG modulations which predict future choice behaviour, we applied conjunction map analyses (Nichols et al., 2005) to investigate coactivation of the decoding topography and regression weight topography from eegGLM 3, which represent RPE, magnitude, and visual surprise (see Supplementary Figure 8 for more information on this model). We found that both, RPE and magnitude, contribute to the decoding topography. In contrast, there was no significant covariation that survived correction for multiple comparison when we included visual surprise to the conjunction map analyses. These results mirror our behavioural findings, where both previous outcome and magnitude influenced participants’ choices. Moreover, these results corroborate previous findings suggesting the P3b as a final path of feedback processing guiding future decisions (Fischer & Ullsperger, 2013; Nassar et al., 2019).

**Figure 6.**
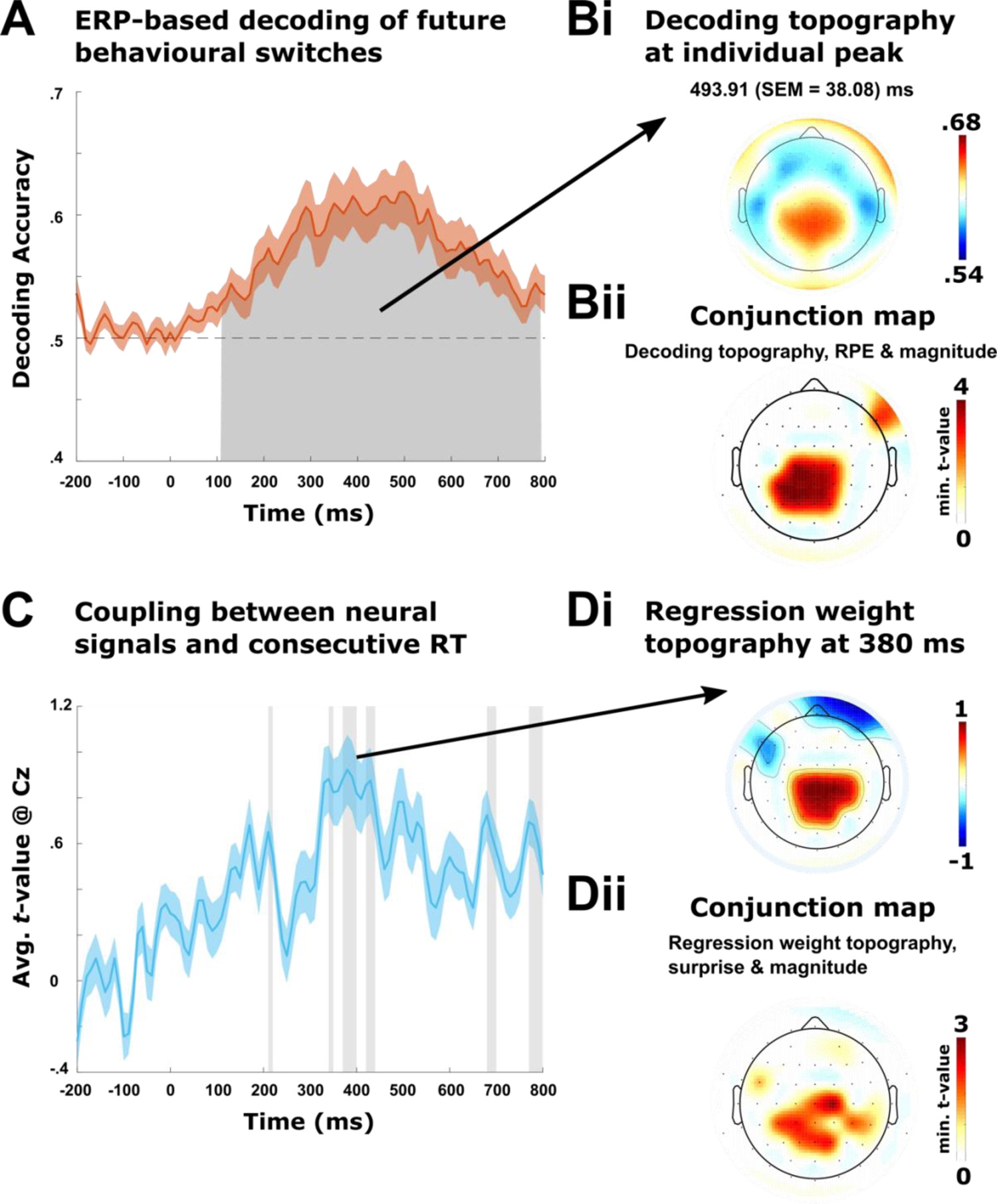
Feedback-related EEG dynamics predict future behaviour. **(A)** Mean accuracy of ERP-based decoding of behavioural switches. Chance-level performance is indicated by the dashed horizontal line. Shadings indicate ±1 SEM. Gray area indicates the cluster of time points in which decoding accuracy significantly surpassed chance level after correction for multiple comparisons using cluster-based permutations analysis (Bae & Luck, 2019; Maris & Oostenveld, 2007). **(Bi)** A search light analysis at the latency of the maximal individual decoding accuracy revealed a centroparietal topography matching the P3b. (**Bii**) Temporospatial conjunction map analyses (Nichols et al., 2005). Minimum *t*-statistics of significant coactivation of all included regressors (decoding accuracy, reward prediction error (RPE) and magnitude) after FDR correction are shown. Results indicate that both, RPE and magnitude, contribute to the decoding topography. *Note*: One participant was excluded from this analysis as there were not enough switch trials for this participant to generate two valid ERPs per category for the test set. (**C,D**) Coupling between neural signals and consecutive RT slowing at Cz (electrode with the maximal effect). Robust regression coefficients indicate across all subjects that the neural signal on a given trial covaries with the following trial’s RT with a peak in the P3a time range. Thus, longer RT’s in the next trial are associated with higher P3a amplitudes on a given trial. Shades represent SEM and the grey shaded areas mark the time of significant effects that survived FDR correction. (**Di**) The topography plot display regression weights at the RT prediction peak and all non-significant electrodes after FDR correction are masked out. (**Dii**) Conjunction map analyses (Nichols et al., 2005) indicating that both the surprise and magnitude regression show a coactivation with the topography of the peak RT prediction.

Additionally, we sought to investigate possible relationships between single-trial feedback-related EEG activity to adaptations in RT in subsequent trials. Therefore, we regressed feedback-related EEG activity at each data point onto RT in consecutive trials including factors of no interest (jitter, trial number). We found a positive covariation between single-trial EEG activity and consecutive RT in the P3a time range displaying a typical, yet slightly more parietal, scalp topography (Cz peak at 380 ms, *t*(23) = 2.98, p = 0.006, *d* = 0.62, CI [.28, 1.56], see Figure 6C and 6Di). Conjunction map analysis (Nichols et al., 2005) revealed a significant coactivation of the surprise and magnitude regressor of eegGLM3 at 380 ms with this scalp topography (see Figure 6Dii), suggesting that neural correlates of the non-normative factors reflecting unexpectedness contribute to RT slowing in consecutive trials.

## Discussion

In this study we examined whether and how factors that are irrelevant for goal achievement (i.e., non-normative factors) alter learning, and which temporospatial EEG correlates accompany these potential biases. We used a reversal learning variant of a previously established task (Fischer & Ullsperger, 2013) that allowed us to investigate contributions of two non-normative factors (i.e., random pay-out magnitudes and sensory surprise) to human participants’ task performance. Consistent with previous work, we found that participants’ learning was influenced by non-normative factors (Kao et al., 2020; McGuire et al., 2014). In line with models of reinforcement learning (Sutton & Barto, 2018), participants updated decision-guiding-stimulus-values based on RPEs. Importantly, we show that the size (via random pay-out-magnitudes) and the impact of RPEs (via visual surprise) on this value-update was inappropriately scaled by non-normative task factors. On the behavioural level, these biases manifest themselves by post-feedback slowing (PFS) on the immediately following trial, and in choice behaviour at the next encounter of the same stimulus, which could occur within the next ten trials. On the neural level, we expected these parameters (RPE and non-normative factors) to be linked to dissociable neural correlates, but also to converge in a final neural pathway guiding future behaviour (Fischer & Ullsperger, 2013). This is exactly what we found. Common to all parameters is that they modulate the stereotypical ERP sequence after visual feedback (i.e., the early visual P1/N1 complex, and the subsequent FRN, P3a/P3b complex (Kappenman & Luck, 2012; Ullsperger, Fischer, et al., 2014)) in some way. Yet, they differ in terms of their temporal dynamics and dependence on feedback factuality. In the following we begin by describing contributions of each parameter to this uniform sequence of EEG activity. We then argue that the central to centroparietal EEG coactivation of all parameters reflects a common final pathway of feedback processing driving future behaviour and learning.

### Early visual EEG activity following feedback

The first significant covariation of feedback-locked EEG activity was found for the visual surprise regressor during both processing of real and fictive feedback. Specifically, we found an early frontocentral negative covariation combined with a positive bilateral occipital covariation around 100ms that was followed by a positive parieto-occipital covariation with polarity reversal at Oz around 200 ms. We interpret this activation as response of the visual system to the surprising background colour change during feedback presentation on 20% of the trials. Indeed, early sensory EEG responses have been shown to be affected by surprise (e.g., Sowman et al., 2012). On the other hand, the frontocentral negative covariation around 100ms post feedback might not just be a mere projection of the effect of the visual system. It could also be, that this covariation already reflects a very early cognitive evaluation of surprise. Supporting this argument, there is evidence of involvement of early activation of the posterior medial frontal cortex (pMFC) during decision making (Mulert et al., 2008). The pMFC has been consistently implicated in performance monitoring, cognitive control, and decision making (Kolling et al., 2018; Shenhav et al., 2016; Ullsperger, Danielmeier, et al., 2014).

Replicating previous findings (Fischer & Ullsperger, 2013), we found a negative early occipital reward prediction error (RPE) effect that occurred around 200ms exclusively after fictive feedback. This effect is likely generated in extrastriate visual areas and parietal posteromedial cortex (Boorman et al., 2011; Fischer & Ullsperger, 2013) and may reflect a mechanism that eases counterfactual learning (Gold & Shadlen, 2007).

### Dissociable contributions to a uniform sequence of EEG activity associated with performance monitoring

A uniform sequence of EEG activity associated with performance monitoring is typically found after feedback: an early frontocentral negativity (the FRN), followed by a frontocentral positivity (the P3a ERP component), that is succeeded by a more sustained parietal positivity called P3b (Ullsperger, Fischer, et al., 2014). In the following, we discuss dissociable contributions of our task factors to this sequence.

The FRN peaks 200 – 300ms after feedback, is modulated by RPEs and surprise, and consistently localized to the pMFC (Ullsperger, Fischer, et al., 2014; Walsh & Anderson, 2012). In line with previous work, we found that the FRN tracks RPE signals (Sambrook & Goslin, 2015), but only if rewards were actually obtained, rather than counterfactual (Fischer & Ullsperger, 2013). This suggests that the FRN may require active involvement to track RPE signals. Supporting this argument, there is evidence that the RPE signal contributing to the FRN is also absent in observational learning (Burnside et al., 2019). An alternative explanation may be a general disengagement of attention during counterfactual or observed feedback to others’ actions. However, this is unlikely given that our participants learn equally well from factual and counterfactual outcomes and that EEG representations of visual surprise in the FRN time window are similar for both conditions. These results suggest that visual surprise is coded in the FRN independent of feedback valance and factuality. This data fits well with the idea that the FRN reflects reward predictions errors for both positive and negative feedback (Sambrook & Goslin, 2015) and is in line with recent research suggesting novelty to affect feedback processing in the FRN time window (Ernst & Steinhauser, 2020). However, the manipulation of the FRN through surprise does not appear to habituate over time (see Supplementary Figure 8D). Thus, visual surprise may alternatively introduce some modulation of the FRN time window through visual-evoked responses. With respect to random pay-out magnitudes, we found that they are also tracked in the FRN during both, real and fictive feedback. This suggests objective coding of the reward magnitude in the FRN, regardless of whether the reward is actually obtained or not. Coding of reward magnitude in the FRN is at odds with a previous study suggesting that the FRN is insensitive to reward magnitudes (Yeung & Sanfey, 2004). In this study, high and low reward magnitudes could be predicted because the particular card colours were consistently associated with high or low reward magnitudes. In contrast, in the present study reward-magnitudes varied randomly and could therefore not be learnt. One possible characterisation of these findings is that under maximal expected uncertainty the FRN tracks reward magnitudes in a similar fashion as surprise (i.e., unexpected high or low reward magnitude), while this is not the case when prior reward magnitude prediction is possible (Yeung & Sanfey, 2004).

Following the FRN, the pronounced midlatency frontal-central effect for real RPEs and feedback-factuality-independent visual surprise fits well with research linking the P3a to attentional allocation of resources to stimuli (Polich, 2007). Our data suggests that negative RPEs trigger attentional orienting and that this is reflected in the increased amplitude of the P3a. However, in line with previous work, this effect appears to depend on whether or not feedback had factual consequences (Fischer & Ullsperger, 2013). In contrast to RPEs, increased feedback salience by visual surprise triggered attention orienting regardless of whether the feedback was factual or counterfactual. This was indexed by a large frontocentral P3a effect. This feedback-factuality-independent effect of visual surprise may be an index of the central nervous response to perceptual novelty (Friedman et al., 2001; Wessel & Aron, 2017). Given that the coloured background of the feedback occurs at a stable frequency of 20%, participants build up an expectation of these rare feedback events, which, in turn, should reduce the surprise elicited by them. Therefore, a reduction (habituation) of surprise-related effects with increasing time-on-task is expected (on a behavioural level, this is reflected in a reduction of post-surprise slowing over time (see Supplementary Figure 9)). In contrast, early visual evoked potentials generated in primary visual cortex should not be reduced over time. We found an interaction of surprise x trial number (see Supplementary Figure 8D) indicating a habituation of the frontocentral surprise effect with increasing exposure to the rare visually surprising feedbacks. This supports our interpretation that this frontocentral P3a-like effect at 300 ms indeed reflects surprise.

As feedback processing continues, the different task factors appear to converge on a common late central parietal positivity that coincides with the P3b ERP component (Polich, 2007). The P3b has been linked to value encoding in working memory and more generally to value updating in the reinforcement learning context (Fischer & Ullsperger, 2013; Polich, 2007). Thus, the effect of RPEs and random pay-out magnitudes on the P3b likely reflect updating and (re-)encoding of a stimulus’ expected value that guides future behaviour independently of feedback factuality. This assumption is in line with more recent work suggesting that the P3b relates to learning (Jepma et al., 2018; Jepma et al., 2016). Nassar et al (Nassar et al., 2019) further disentangled the relationship between learning and the P3b. They demonstrated that the P3b does not naively reflect increased value-updating but may adaptively adjust learning depending on the statistical context of the observed surprising event (i.e., increasing learning from surprising events indicative of chance points and decreasing learning from uninformative oddballs). In the present study, we do not see a strong modulation of the P3b though visual surprise. One explanation for this effect might be that participants quickly learn that visual surprise is uninformative for stimuli values. Therefore, the effect of visual surprise may be reflected during earlier feedback processing (see above) and its processing may be terminated before influencing value update and future choice behaviour. In contrast, the effect of oddballs on belief updating in the Nassar et al (2019) study may have to be actively suppressed via P3b activity.

### A common late centroparietal positivity serves as a final pathway of adaptation that guides future behaviour

Taken together, the results discussed above suggest that the FRN/P3a/P3b complex is dynamically and in a spatio-temporally dissociable fashion driven by representations of multiple outcome parameters deemed relevant by the learning agent. These are RPEs and other motivationally relevant, yet non-normative, parameters like simple visual surprise and random pay-out magnitudes. While the FRN and P3a likely require active involvement to track RPEs signals, other motivationally salient parameters are tracked in this ERP complex irrespectively of feedback factuality. Later in feedback processing these streams of processing decision-related parameters (including subcortical processing of RPEs for counterfactual feedback, cf. Jocham et al. (2014)) converge on a common integrative process reflected in a centroparietal positivity. In line with previous research (Chase et al., 2011; Fromer et al., 2021; Nassar et al., 2019; Ullsperger, Fischer, et al., 2014; Wessel & Aron, 2017) we suggest, that this EEG correlate reflects a final pathway of adaptation that sets the stage for future behaviour. This view is supported by our decoding analyses suggesting that behaviour switches are best predicted by an EEG activity with a topography and latency matching the P3b. Regression weight topographies of the feedback-locked RPE and magnitude regressor contributed to this decoder’s topography. These results fit well with our behavioural and modelling results, suggesting that participants’ choices are affected by previous outcomes, pay-out magnitudes and behaviour. In line with another prominent theory linking the P300 to the anticipation of the need to inhibit responding after unexpected events (Wessel, 2018; Wessel & Aron, 2017), we were able to predict post-feedback slowing on the immediately following trial with a centroparietal topography matching the P3b. Interestingly, the peak of this prediction preceded the peak prediction of choice switches by approximately 100ms and was driven by both visual surprise and random pay-out-magnitudes. This may fit well with the idea that cognitive shifts after unexpected events are preceded by motor inhibition signals (Wessel & Aron, 2017) and suggest that the latency of the centroparietal positivity plays a role in pinning down different aspects of future behaviour.

When discussing the neural representations of task factors relevant for future behaviour, it is worth mentioning that their representation is broader than the centroparietal positivity most predictive for future adaptions. For example, we show that our parameters also modulate more frontal parts of the FRN/P3a complex, with larger effects sizes. However, our data suggest that the centroparietal positivity coactivation seems to best predict future behaviour. Moreover, the FRN/P3a complex does not have to be elicited to initiate this final pathway of adaption. This is shown by the absence of the RPE effect in the FRN/P3a time window after counterfactual feedback. Here RPEs seem to be coded solely subcortically, specifically in the striatum (Jocham et al., 2014), whose activity is not projected to the scalp-recorded EEG.

## Conclusions

We found that individuals are unable to ignore non-normative factors during decision making despite explicit knowledge of the irrelevance to these factors for optimal task performance. Specifically, uninformative reward magnitudes alterations and sensory surprise biased learning behaviour in a novel probabilistic choice task. On the neural level, these parameters are represented in a dynamic and spatiotemporally dissociable sequence of EEG activity. Later in feedback processing the different streams converge on a P3b-like central to centroparietal positivity that serves as a final pathway of adaptation and governs future behaviour.

## Materials and Methods

### Participants

Twenty-eight healthy participants were recruited into this study. The data of four participants were excluded because of technical problems (n = 1) or because they performed the task at chance level (n = 3). The final sample consisted of 24 participants (18 female, mean age: 24.25 (SD = 4.12)). All participants were informed about the experimental procedures and gave written, informed consent. The study protocol was approved by the local ethics committee.

### Experimental task

To maximize financial earnings in this probabilistic choice task, participants had to learn the changing reward probabilities of three different stimuli. At each trial, participants could either decide to gamble on a specific stimulus and win or lose 10 or 80 points (translating to 0.10/ .80 EUR) or choose to avoid the stimulus and observe what would have happened, without any financial consequences. The task was administered using Presentation 16.3 (Neurobehavioral Systems).

The task consisted of seven blocks in which the three stimuli were alternated but their reward probability was kept constant. In each block, stimuli were presented for at least 26 times and not more than 42 times. In total, the task comprised 18 reward probability reversals and 720 trials. Reward probabilities could either be low (20%), neutral (50%), or high (80%). Importantly, we introduced two types of unexpected events that were irrelevant for optimal task performance but nevertheless could potentially bias learning: a) simple visual surprise, where on 20% of the trials, the standard colour of the feedback background (black) was briefly replaced by a colour matching the feedback colour (i.e., background flash in green for positive feedback and in red for negative feedback, respectively; see Figure 1Aii), and b) random pay-out magnitudes. Here, reward magnitudes randomly varied, so that on 50% of the trials 10 points and on the other half of the trials 80 points could be won or lost (see Figure 1Aiii).

Each trial began with a random jitter between 300 ms and 700 ms, during this jitter period a central fixation cross and two response options (choose – green tick mark or avoid – red no-parking sign) were shown (see Figure 1Ai). The response options sides were counter-balanced across participants and remained in place until the feedback was presented. After the fixation cross, the stimulus was shown centrally until the participant responded or for a maximum duration of 1700 ms. If participants failed to respond in time, they were instructed to speed up. Thereafter, participants’ choices were confirmed by a white rectangle surrounding the chosen option for 350 ms. Finally, the outcome was presented for 750 ms. If subjects chose to gamble on the presented stimuli, they received either a green smiling face and a reward of 10 or 80 points or a red frowning face and a loss of either 10 or 80 points. When subjects avoided a symbol, they received the same feedback but with a slightly paler color and the points that could have been received were crossed out to indicate that the feedback was fictive and had no monetary effect. During visual surprise trials, the feedback was presented on a feedback-matching colour (green or red; see Figure 1Aii).

### Behavioural regression analyses

We performed multiple-robust regression analyses (Fischer et al., 2018), predicting either participants decisions to gamble or avoid a stimulus or their reaction times. We performed regressions on two subsets of trials: First we run the regression on the original trial order. Here, stimuli appeared in intermixed fashion allowing us to investigate the influence of directly preceding trials on participants behaviour on a given trial (bGLM 1 & 2). Second, to examine how the behaviour and task factors on the last encounter of a specific stimuli influenced the behaviour on the next encounter of the same stimulus, we resorted the trial order according to stimulus identify (bGLM 3 & 4).

The logistic choice model in the original trial order was given by:

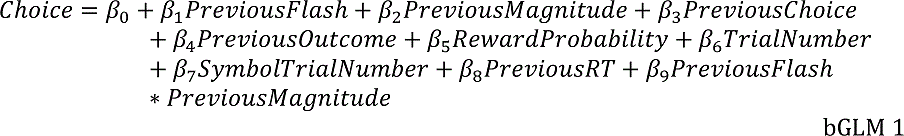

The logistic choice model run in resorted trial order was defined by:

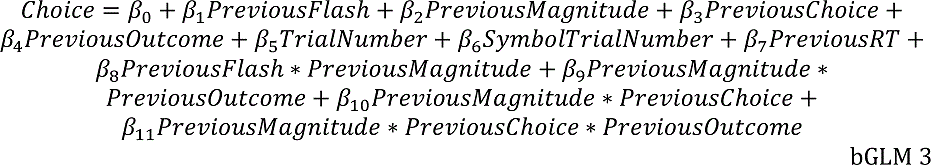

The regression model on RT in the original trail order included the following regressors:

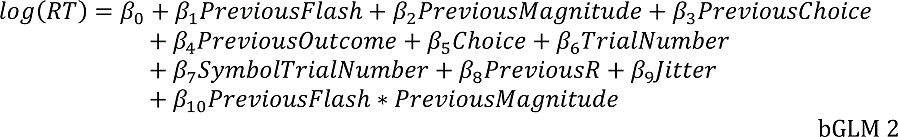

The regression model on RT in resorted trail order is described below:

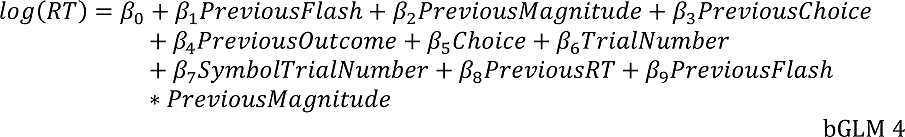

The individual factors are: Previous Flash = surprise in the previous trial (bGLM1 & 2) / last stimulus encounter (bGLM3 & 4): 0 = no visual surprise, 1 = visual surprise). Previous Magnitude = low (0) or high (1) reward magnitude in the previous trial (bGLM1 & 2) / last stimulus encounter (bGLM3 & 4). Previous Choice: coded participants choice (0 = avoided, 1 = chosen) in either the last trail (bGLM1 & 2) or last encounter of the same stimuli (bGLM3 & 4). Previous Outcome = outcome of the immediately preceding trials or stimulus encounter (0 = loss, 1 = win). Reward Probability = reward probability of the current stimuli. Trial Number = log-scaled trial number (reflecting the time in the task). Symbol Trial Number = block-trial number for each symbol (reflecting how often the symbol has been seen in the respective block). Trial number and symbol trial number served mainly to control for unspecific effect of task duration, like fatigue. Regression analyses were performed separately for each participant. Individual participants t-values per regressor were then tested on group level via two-sided *t*-test against zero (*p*-values were corrected for multiple comparisons (0.05/number of regressors)). See supplementary materials for a detailed visualization of all the results of the bGLMs.

### Computational modelling

We fit 4 different models to participants’ choice data:

**Model 1**: Standard model. This was a standard Q-learning model. Here, the expected value of an action was calculated as follows:

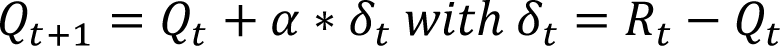

Q values represent the expected value of an action at trial t. α reflects the learning rate (i.e. to what extend does new info override old info). δ_t_ represents the prediction error with *R_t_* being the reward magnitude of that trial. In this model, high reward magnitudes were weighted eight times higher than low magnitudes. The likelihood for the model to choose a stimulus was calculated according to a softmax rule:

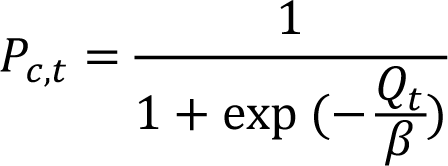

The free parameter β reflects choice stochasticity, with high values corresponding to more random choices and low values corresponding to more deterministic choices.

**Model 2**: Binary model. This model is identical to model 1, except that high and low reward magnitudes are scaled equally (namely all wins = 1 and all losses = 0).

**Model 3.** Outcome-weighted model. This model is identical to model 1, with the addition of an outcome weighting parameter γ, that can down-weight or up-weight outcome:

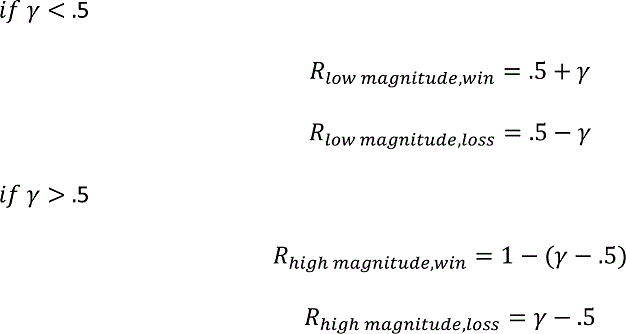

Specifically, we first scaled all rewards equally (all wins = 1 and all loss = 0). Second, depending on its values the free parameter γ could either up-weight low outcome magnitudes (γ< .5) or down-weight high outcome magnitudes (γ> .5). Hence, γ regulates the effect of the reward prediction error on value update, as described in Model 1:

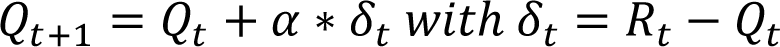

**Model 4.** Outcome weighted plus surprise model. This model is identical to model 3, except that α can change on visual surprise trials through the parameter λ:

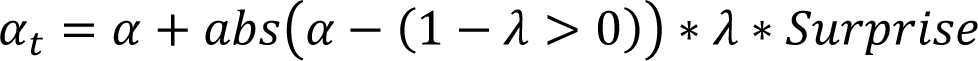

The second term in this equation calculates the distance to maximal (1) or minimal (0) learning rate depending on parameter λ (i.e., for a positive λ this term is the absolute difference between alpha and the maximal learning rate 1; for negative λ this term is simply the learning rate α). The term Surprise is a Boolean expression that is true (i.e. 1) for trials with visual surprise and false (i.e. 0) for trials without surprise. Here, λ can vary between −1 (i.e., learning rate becomes 0 on visual surprise trials) and 1 (then the learning rate becomes 1 on visual surprise trials). In general, positive λ values suggest increased learning and thereby up-weighting of the effect of the reward prediction error on value update on respective trials. Negative values have the opposite effect, i.e., reflect a distracting effect of the visual surprise.

### Parameter estimation and model comparison

Parameters were optimized using custom-written scripts in MATLAB R2017a (The Mathworks Company, Natick, MA) and constrained optimization using MATLAB’s function fmincon to minimize the sum of the negative log likelihood of the data given the parameters and negative prior probability (Gershman, 2016). See supplementary materials for empirical prior distribution and hyperparameters for parameter constrains. The integrated Bayesian Information Criterion (iBIC) was used to compare between different models (Huys et al., 2011). Here, smaller values indicate better and more parsimonious model fits. To validate our models, we applied model and parameter recovery analyses (Wilson & Collins, 2019), reported in the supplementary materials (see Supplementary Figure 1).

### EEG measurements and analyses

Scalp voltages were recorded and A-D converted from 64 channels using BrainAmp MR+ amplifiers (Brain Products, Gilching, Germany) at a sampling rate of 500 Hz. The ground electrode was located at AFz. Data was recorded with CPz as reference channel and re-referenced to common average for analyses. Pre-processing of the EEG data was done under Matlab 2017b (The MathWorks, Natick, MA) and the EEGlab 13 toolbox (Delorme and Makeig, 2004) using custom routines and following pre-processing steps described previously (Kirschner et al., 2020). Specifically, pre-processing steps included: (1) Filtering (0.2 Hz high- and 40 Hz low-pass filter), 2) re-referencing to common average, 3) Segmentation into feedback-locked epochs spanning from 3,500 ms pre-feedback to 1,400 ms post-feedback, 4) automatic epoch rejection, 5) removal of blink and eye-movement components using adaptive mixture independent component analysis (AMICA; Palmer et al., 2012). Following baseline correction (−150 to −50 ms relative to feedback onset), epochs spanning −400 ms to 1000 ms around feedback presentation were then used for multiple robust single-trial regression analyses (Fischer et al., 2016; Fischer & Ullsperger, 2013).

Based on previous findings suggesting differences in feedback processing between real and fictive outcomes (Fischer & Ullsperger, 2013; Schuller et al., 2020), we split up the data according to feedback reality and analysed effects of learning and surprise signals in separate GLMs (real feedback-eegGLM1 & fictive feedback – eegGLM2). These models included single-trial estimates of the signed reward prediction errors (RPE) derived from Model 4 and regressors coding random magnitudes (low vs. high) and visual surprise (no surprise vs. surprise). In addition, both models included trial number and jitter length as regressors of no interest to account for unspecific task effects.

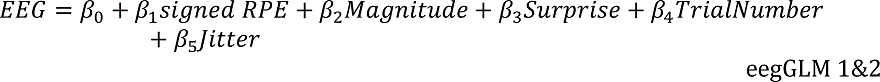

These analyses resulted in regression coefficients for every time point and electrode, revealing the time course and scalp topographies of the relationship between each predictor and neural activity. Resulting p values were corrected for multiple comparisons using false discovery rate (Benjamini & Yekutieli, 2001).

### Multivariate pattern analyses

We used averaged single-trial neural activity (epochs spanning from −200 ms to 800 ms after feedback presentation) of the whole scalp to train a support vector machine to classify whether a participant switched behaviour from choosing to avoiding a gamble or from avoiding to choosing a gamble on the next encounter of the same stimulus identity (on average, 3 trials later (range 1-9)). Following recent recommendations (Bae & Luck, 2018; Grootswagers et al., 2017) we averaged 16 trials per condition (switch vs. no switch). We applied the support vector machine functions implemented in MATLAB 2017b (fitcsvm, predict). The ERP data was smoothed by averaging −10 ms to +10 ms around each datapoint with a step size of 10 ms throughout each epoch. All input data was z-scored across and within electrodes and time. A 100-fold cross-validation using 80% of the trials as training and 20% of the trials (but at least two ERPs per condition) as prediction set was applied. Accuracy was calculated as the percentage of overlap between predicted labels and the ground truth at each datapoint and for each participant. Available trials for each condition were matched with a reduction to the smaller data size via random subsampling. We used cluster-based permutation analyses (Bae & Luck, 2019; Maris & Oostenveld, 2007) to control for multiple-comparison and identify time points of correct classification above chance-level. To localize the information for the classification, we applied a searchlight analysis approach (Fischer et al., 2016) using the same settings as described above. Specifically, we calculated the average accuracy for each electrode alone, every lateralized electrode together with its contralaterally located electrode, and each electrode clustered with the 7 nearest neighbouring electrodes. This resulted in an average accuracy per electrode and time point. To investigate which task factors contribute to the EEG representation of future behavioural switches (i.e., the decoding topographies), we adopted a conjunction map approach (Nichols et al., 2005). Here, we display the minimum *t*-statistics topography against the null hypothesis that one or more effects are null. *t*-statistics for the decoding topographies were derived from *t*-test against chance level and *t*-values for task factors were the result of eegGLM 3. This GLM was defined as followed:

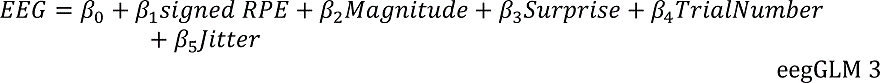

This model comprised both, real and fictive feedback trials. The sign of the RPE was inverted for fictive feedback trials in this analysis (see Supplementary Figure 7 for a summary the results of eegGLM 3). We report temporospatial conjunction maps displaying minimum *t*-statistics of the included factors and mask out all non-significant electrodes after FDR correction (Benjamini & Yekutieli, 2001).

## Acknowledgements

We would like to thank Laura Waite for help with programming the task and collecting EEG and behavioural data.

## Data and code availability

The data of this study can be downloaded on the Open Science Framework at [URL to be inserted after acceptance]. All materials, processing and analysis scripts are available from the authors upon reasonable request.

## Author contributions

A.G.F and M.U. designed the study. H.K. and A.F. analysed the data. H.K. wrote the manuscript. All authors reviewed and edited the manuscript.

## Supplementary Methods

**Supplementary Table 1.**
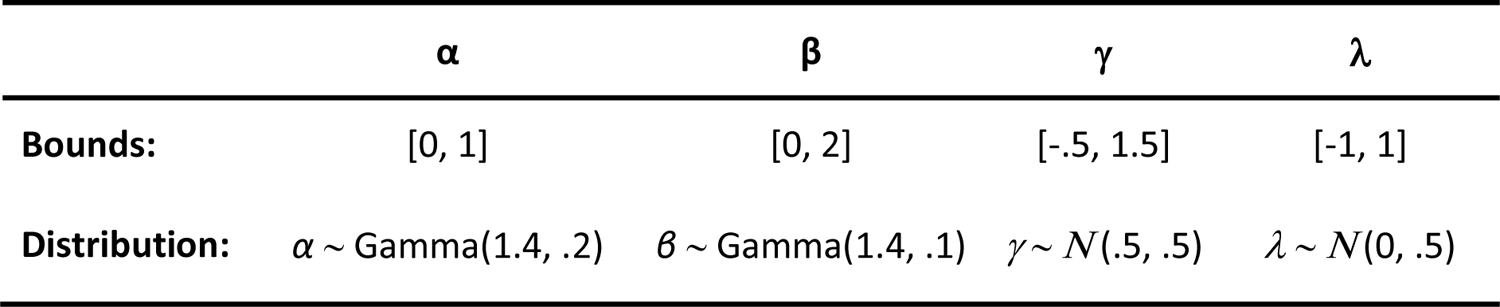
Prior distributions and hyperparameters for parameter estimation. Gamma distributions are parameterized in terms of shape and scale. Normal distribution is parameterized by mu and sigma.

## Supplementary Results

**Supplementary Figure 1.**
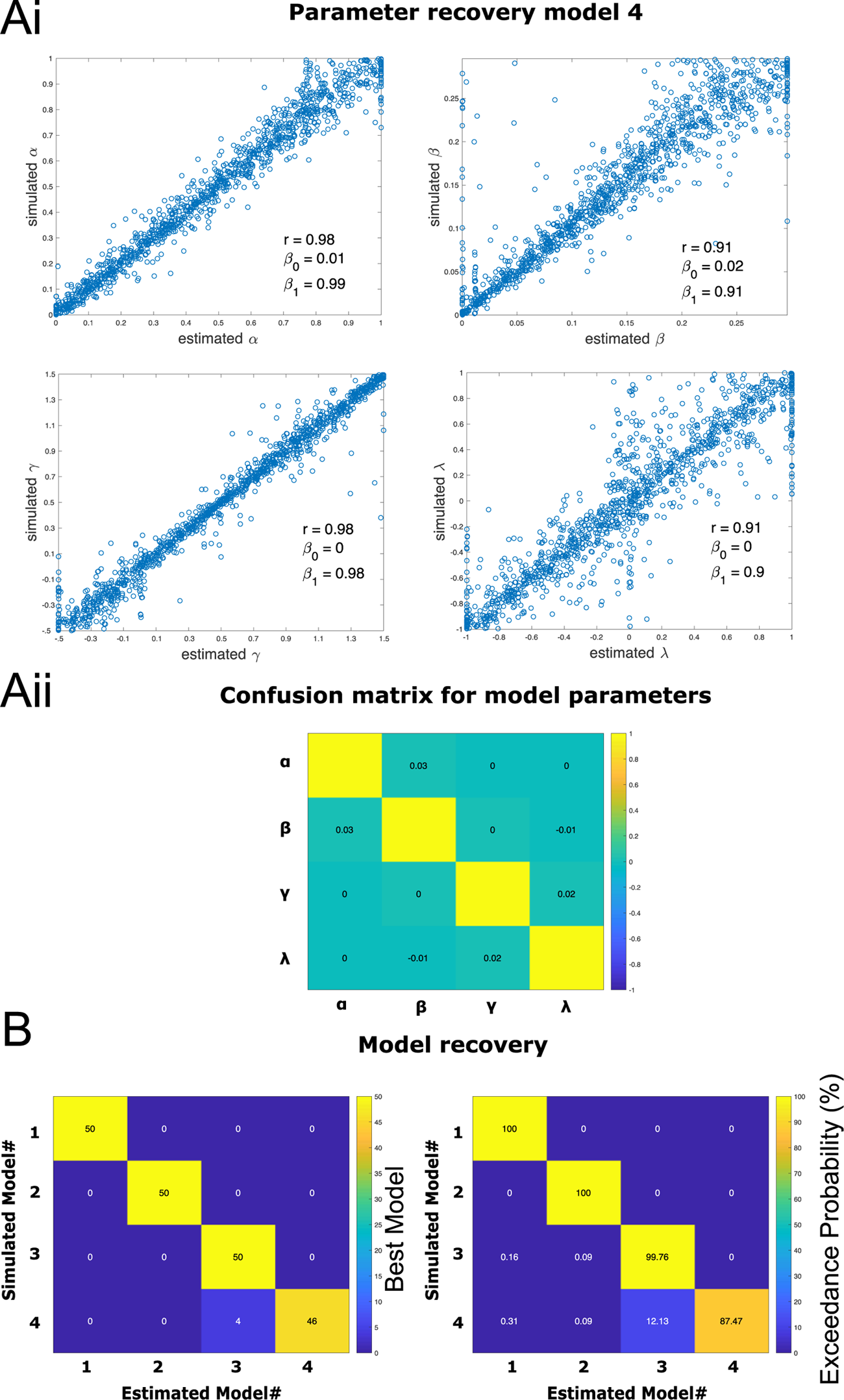
Model and parameter recovery analyses. **(Ai)** Parameter recovery. Data form 1200 surrogate subjects (50 simulations x 24 individuals) was simulated with Model 4 (Outcome weighted plus surprise model). Model free parameters were randomly drawn for each surrogate participant, whereby range for the β-values was limited to the empirically observed range (see Figure 3B). The other parameters covered a wider range than empirically observed. Task properties and contingencies were identical to those of the 24 participants reported in this study. The estimated parameters (Matlab function fmincon) were regressed against the parameters used to simulate the data (i.e. the true parameters). Results show good identifiability with correlation coefficients (r) > .9 and regression intercepts (β_0_) close to 0 and regression slopes (β_1_) close to 1. **(Aii)** The confusion matrix indicates no trade of between parameters of the model. **(B)** Model recovery. For model comparisons of the simulated data (50 simulations x 24 participants x 4 models), the mbbvb-toolbox (http://mbb-team.github.io/VBA-toolbox/) was used to compute the exceedance probability for each model in comparison to all other models. Model free parameters were randomly drawn for each surrogate participant. **(Bi)** All models were robustly identified as the most probable model when they were used to generate the data. **(Bii)** The average exceedance probability was > 85% for all models. Model 1 = standard model; Model 2 = binary model; Model 3 = outcome weighted; Model 4 = outcome weighted plus surprise.

**Supplementary Figure 2.**
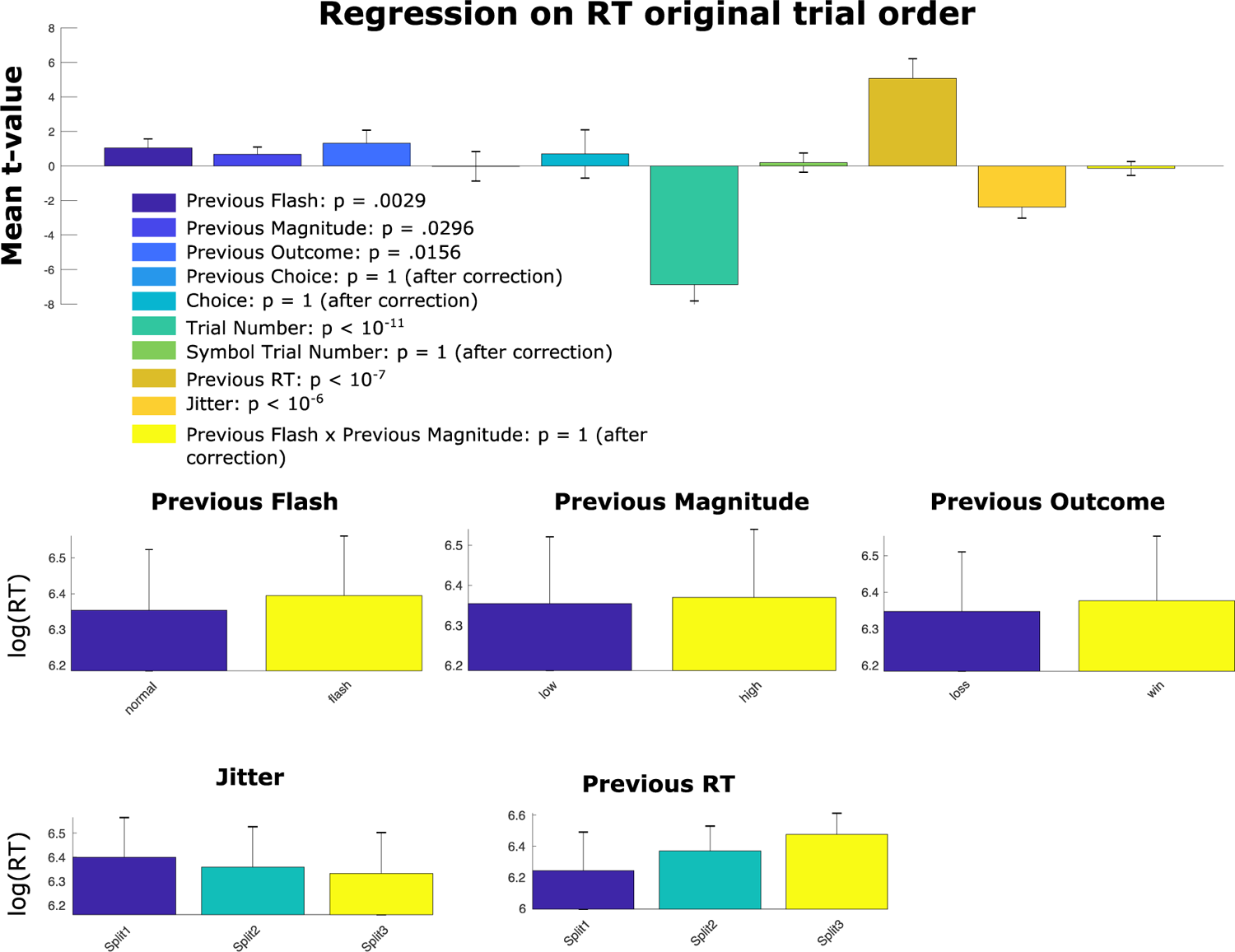
Single-trial regression on log(RT) in original trial order. *Note:* All follow-up plots show raw log RTs, error-bars reflect 99% CI for regression weights, and 99.99% CI for raw data. p-values are derived from t-tests of the individual regression t-values against zero. Results are adjusted for multiple comparisons via Bonferroni correction.

**Supplementary Figure 3.**
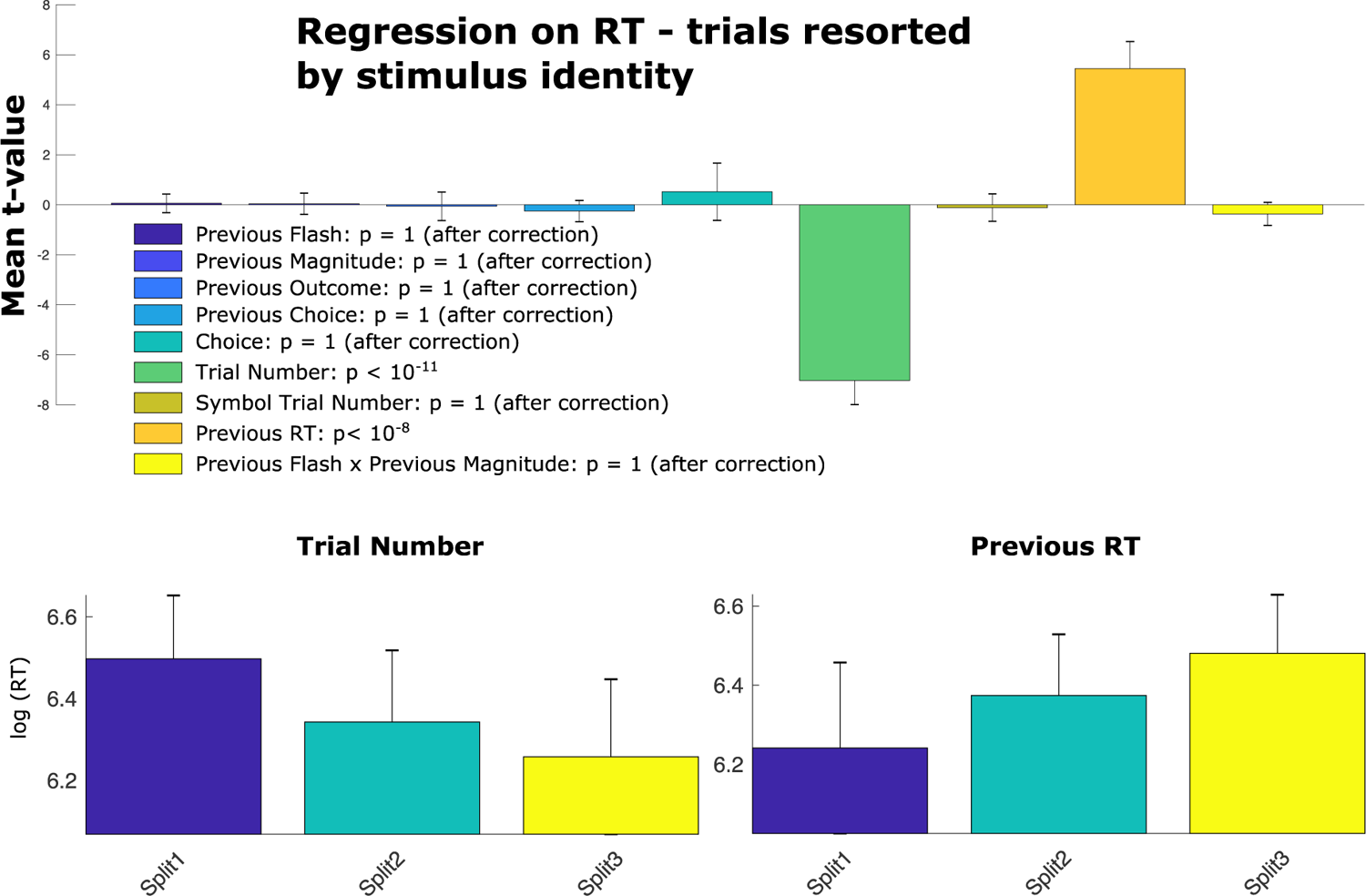
Single-trial regression on log(RT) in original trial order. Multiple single-trial regression on RT. Here, trial order has been resorted according to stimuli identity before running the analysis. Results suggest that participants responded faster as the task progressed and that that the RT during the last stimulus encounter was positively related to the RT in the next time this stimulus was seen. The surprise effects did not affect RT in the next stimulus encounter. *Note:* All follow-up plots show raw log RTs, error-bars reflect 99% CI for regression weights, and 99.99% CI for raw data. p-values are derived from t-tests of the individual regression t-values against zero. Results are adjusted for multiple comparisons via Bonferroni correction.

**Supplementary Figure 4.**
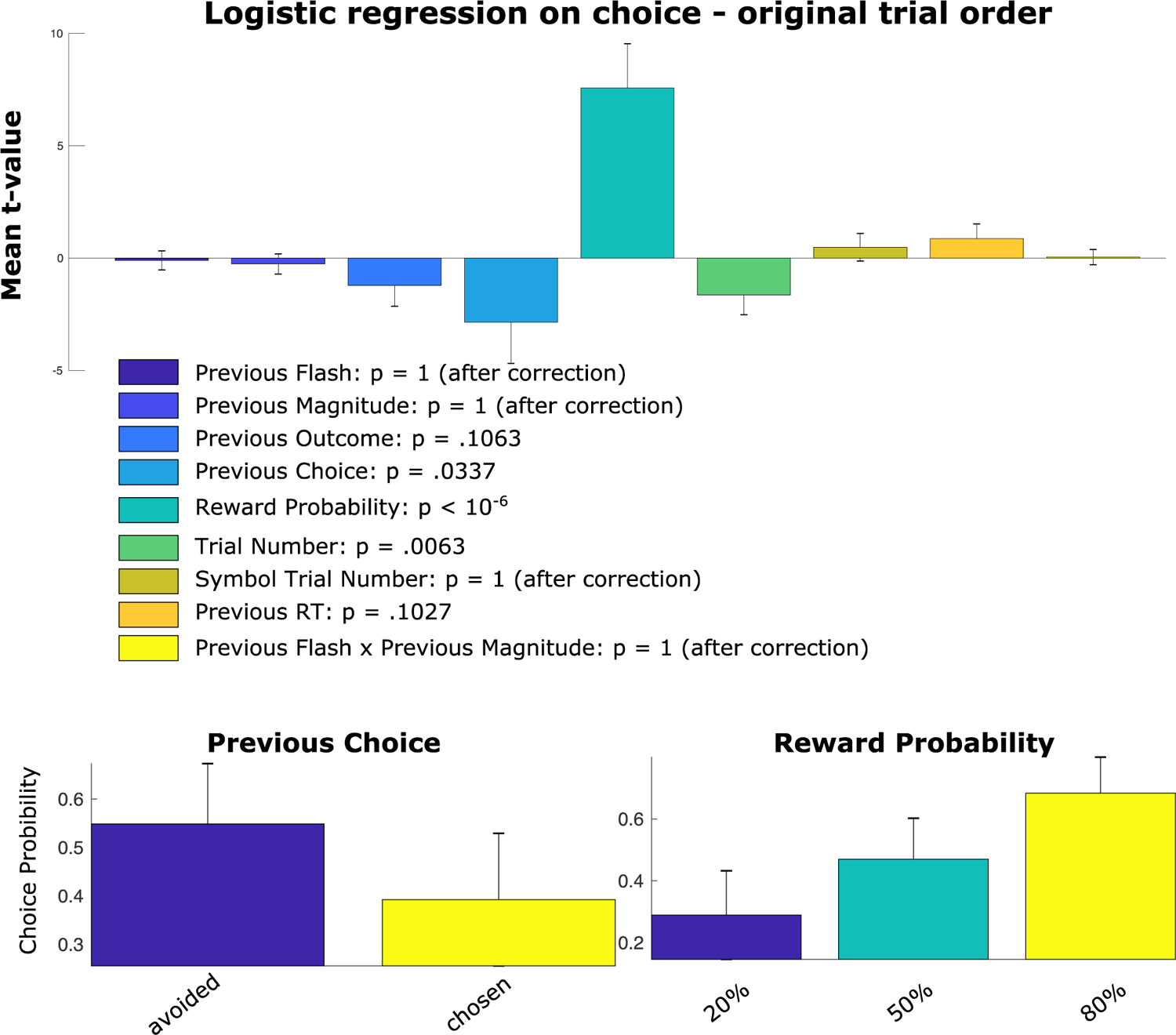
Single-trial logistic regression on choice in original trial order. Multiple single-trial logistic regression on choice behavior. We found that participants were more likely to gamble on a stimulus if the stimulus was previously avoided. However, this choice bias is fairly small and difficult to interpret as on the previous trial participants could have seen both the same stimulus or a different stimulus. The significant reward probability regressor suggests, that participants were able to reflect the reward probabilities of the respective stimulus in their choices. *Note:* All follow-up plots show average choice probability, error-bars reflect 99% CI for regression weights, and 99.99% CI for raw data. p-values are derived from t-tests of the individual regression t-values against zero. Results are adjusted for multiple comparisons via Bonferroni correction.

**Supplementary Figure 5.**
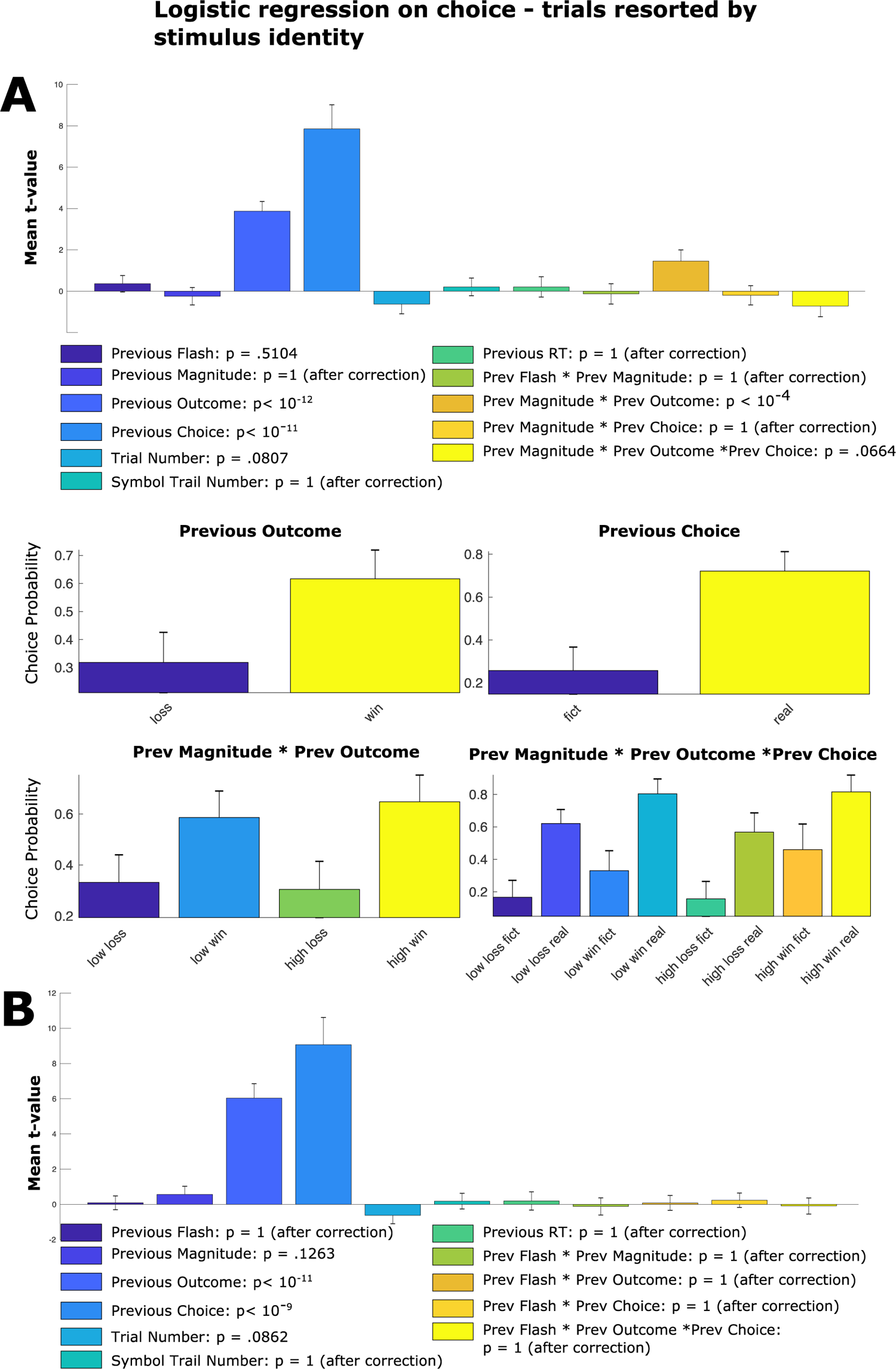
Single-trial logistic regression on choice in resorted trial order. **A)** Multiple single-trial logistic regression on choice behavior. Here, trial order has been resorted according to stimuli identity before running the analysis. We show that participants were more likely to gamble on a stimulus if the stimulus was previously chosen and winning. These effects are modulated by previous reward magnitudes, whereby subjects were more likely to gamble on a stimulus after having received a high counterfactual reward. Moreover, participants were less likely to choose a stimulus after high factual losses. **B)** To investigate if previous surprise had a similar effect on the previous choice and outcome regressor as magnitude, we rerun the regression analyses looking at the interactions for visual surprise. Here, we do not see a similar modulation for visual surprise. *Note:* All follow-up plots show average choice probability, error-bars reflect 99% CI for regression weights, and 99.99% CI for raw data. p-values are derived from t-tests of the individual regression t-values against zero. Results are adjusted for multiple comparisons via Bonferroni correction.

**Supplementary Figure 6.**
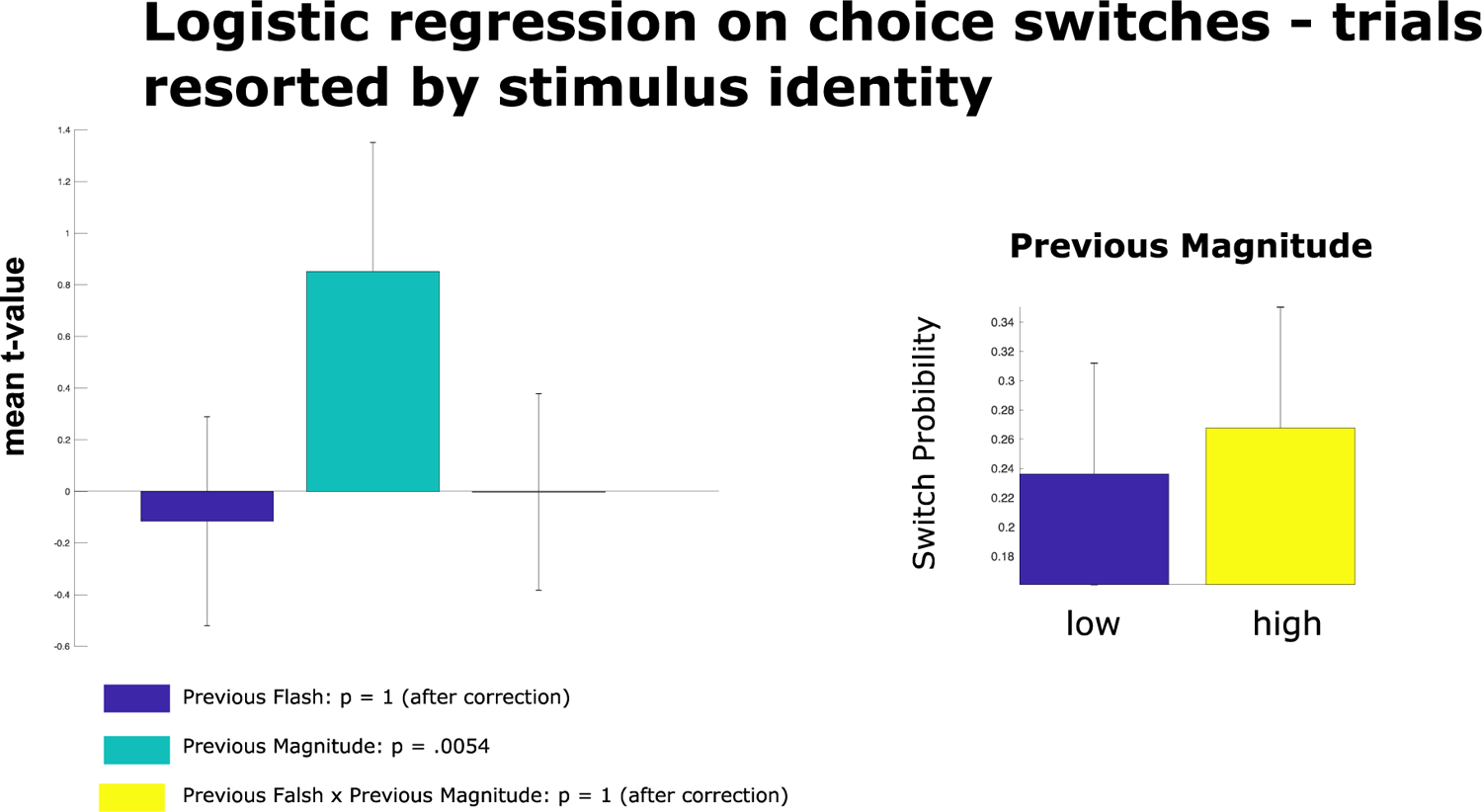
Single-trial logistic regression on behavioral switches in resorted trial order. Here, trial order has been resorted according to stimuli identity before running the analysis. We show that participants were more likely to switch their behavior if the stimulus obtained a high reward magnitude previously. Previous visual surprise did not increase the probability of behavioral switches. *Note:* All follow-up plots show average choice probability, error-bars reflect 99% CI for regression weights, and 99.99% CI for raw data. p-values are derived from t-tests of the individual regression t-values against zero. Results are adjusted for multiple comparisons via Bonferroni correction.

**Supplementary Figure 7.**
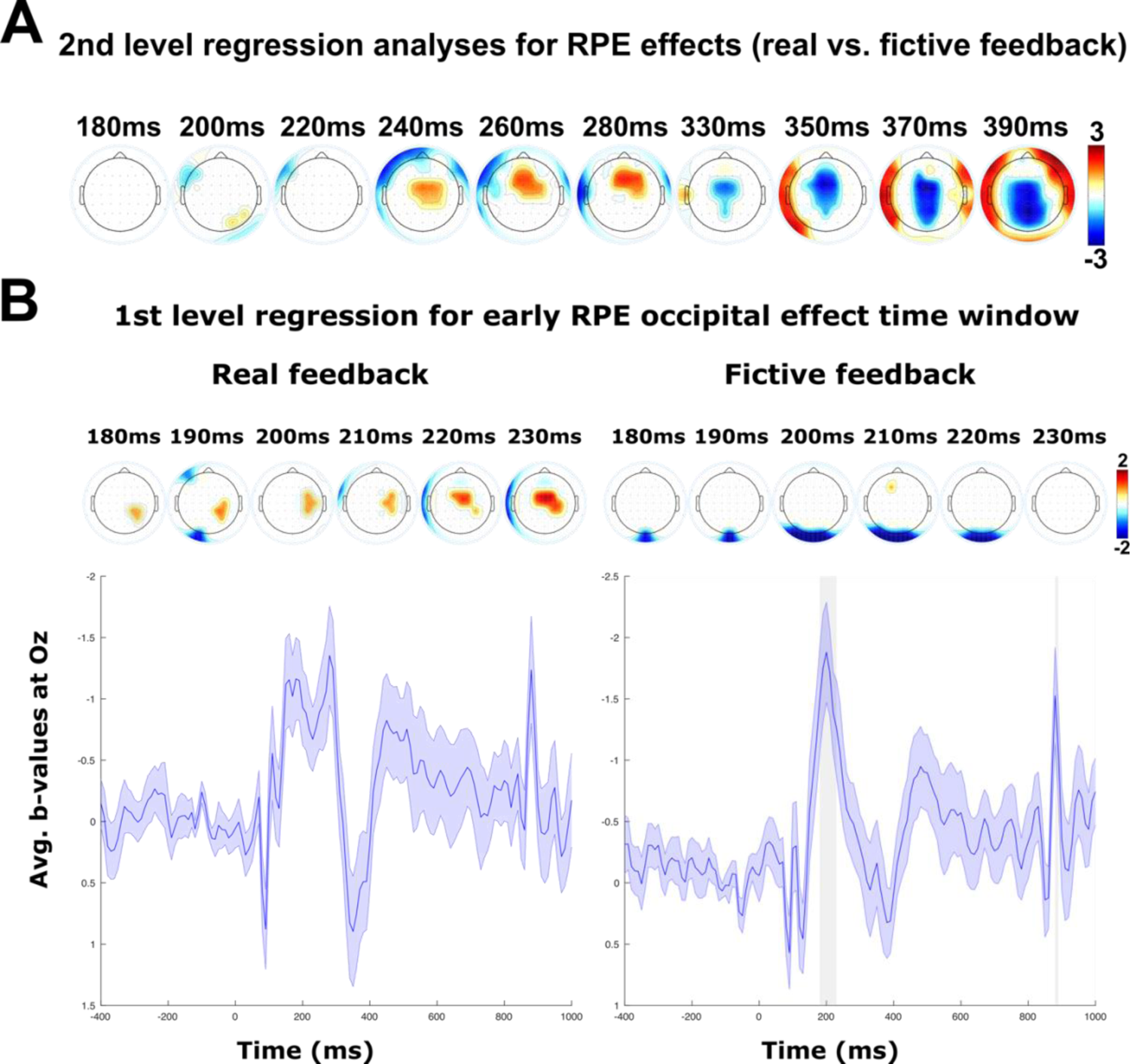
Exclusiveness test of early reward prediction error (RPE) effects. **A)** The topography plots display second level regression weights (real vs. fictive feedback) and all non-significant electrodes after FDR correction are masked out. Results validate the exclusiveness of the FRN and P3a effect for real feedback. However, the exclusive early occipital effect found for fictive feedback did not survive FDR correction. The strong centroparietal effect starting around 350 ms reflects that RPE was calculated with respect to objective stimulus values, whereas the centroparietal EEG is deviated positively when the outcome is subjectively more unfavorable. Objective feedback valence and subjective valence are opposite in fictive outcomes (a missed reward is unfavorable and an avoided loss is favorable) **B)** Follow-up analyses on the exclusiveness of the early occipital effect found predominantly during fictive feedback processing. Topographies show beta coefficients for the RPE regressor of eegGLM1 thresholded at critical p-value from FDR correction. Below mean regression weights of the RPE regressor are shown at Oz. The grey shaded area marks the time of significant effects that survived FDR correction. These results show, that the early occipital effect is significant only in during the processing of fictive feedback.

**Supplementary Figure 8.**
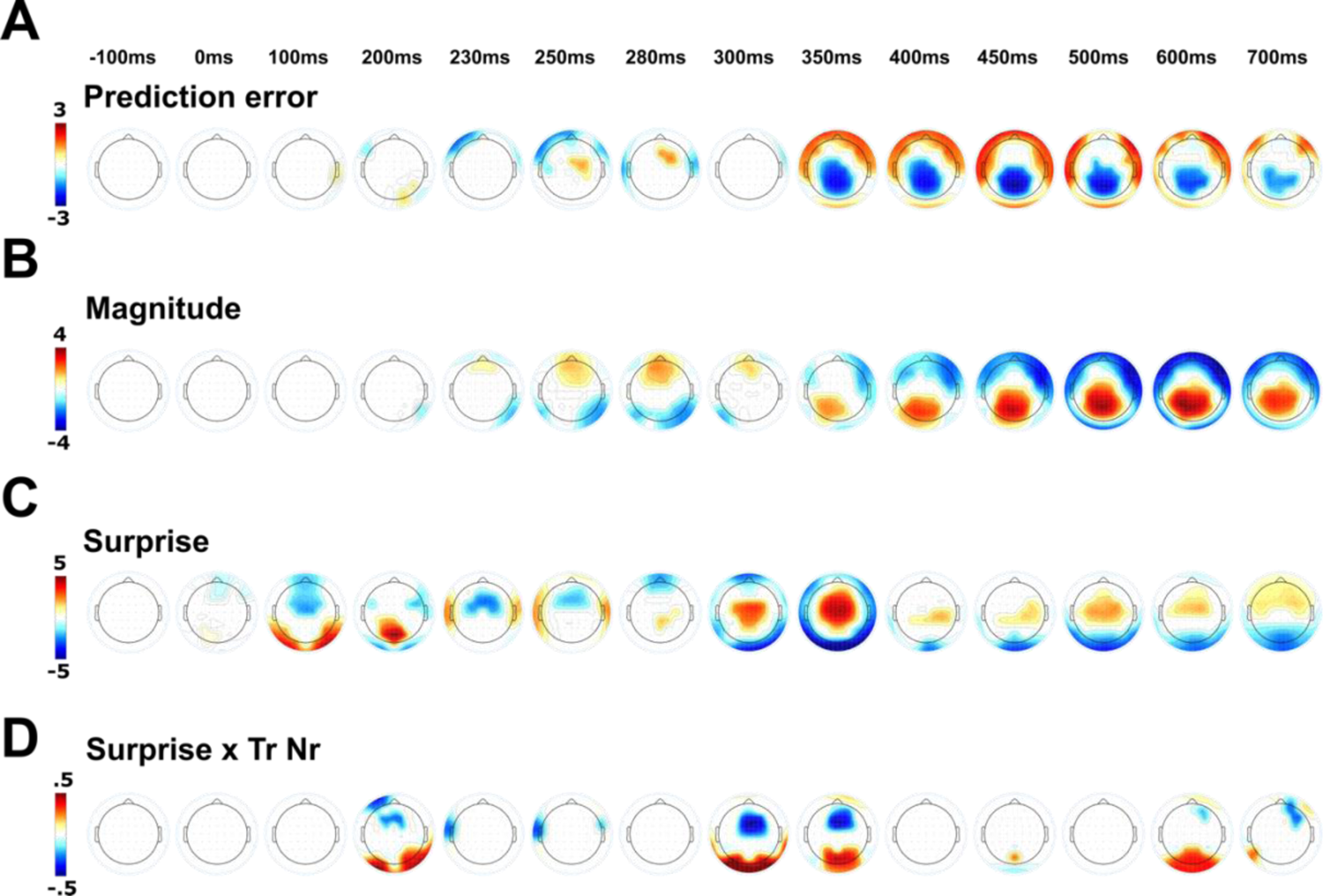
Single-trial regression results of eegGLM3. Time course of regression weight topographies for the feedback-locked reward prediction error. **(A)**, magnitude **(B),** surprise **(C),** and surprise x trial number interaction **(D)** regressors from eegGLM3 to which data from all trials was submitted. Topographies show beta coefficients thresholded at critical p-value from FDR correction. *Note*: In contrast to eegGLM, 1&2, the sign of the RPE was inverted for fictive feedback trials in this analysis. **(D)** significant effects reflect habituation of visual surprise in the EEG activity.

**Supplementary Figure 9.**
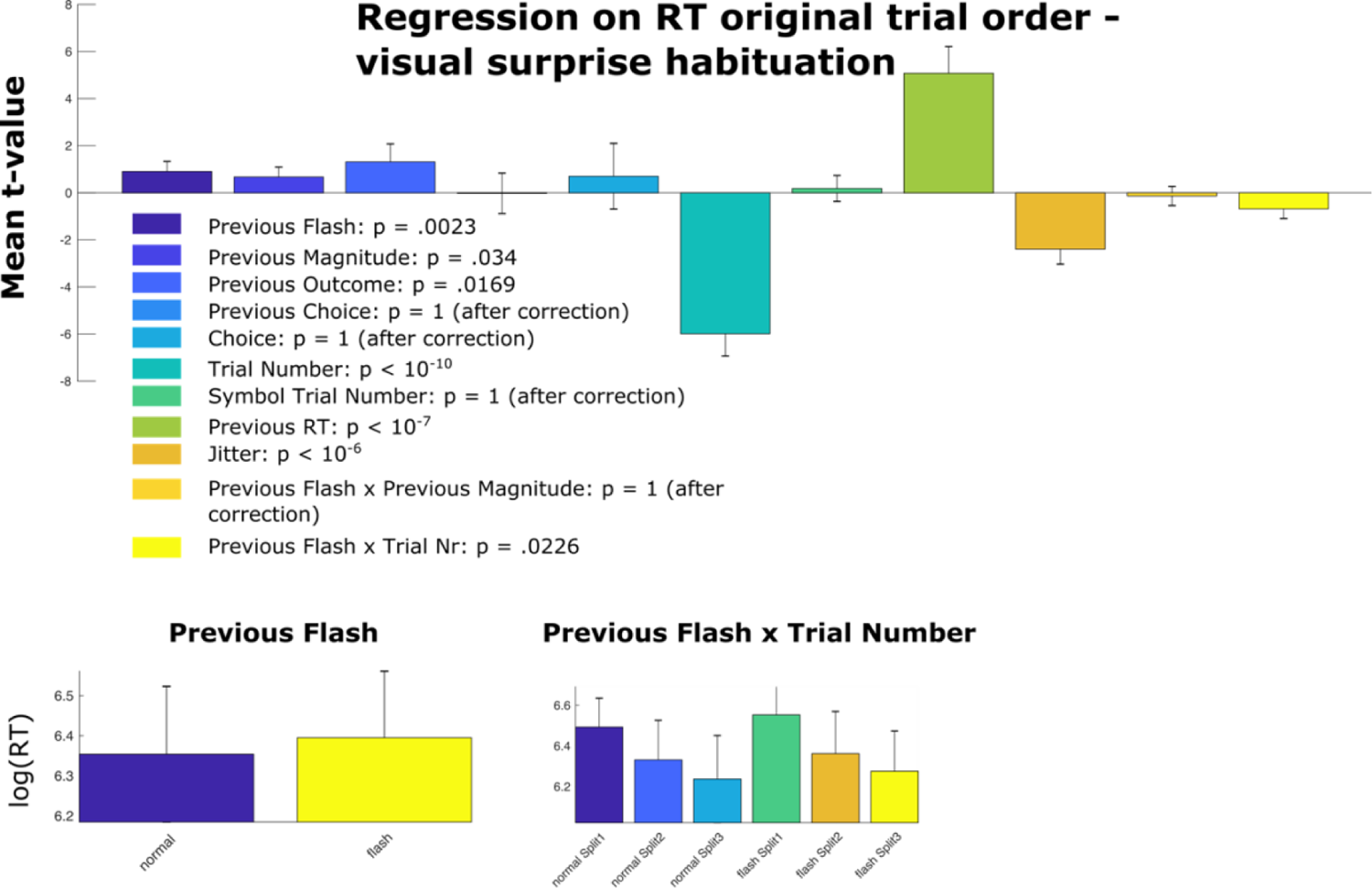
Single-trial regression on log(RT) in original trial order. *Note:* This is bGLM3 extended by the predictor previous flash x trial number to confirm a reduction (habituation) of the post-visual-surprise-slowing (PVSS). We found an interaction between surprise (flash) and trial number indicating a habituation of PVSS. All follow-up plots show raw log RTs, error-bars reflect 99% CI for regression weights, and 99.99% CI for raw data. p-values are derived from t-tests of the individual regression t-values against zero. Results are adjusted for multiple comparisons via Bonferroni correction.

